# Nuclear remodeling and optimal migration emerge early, and constriction-passage progressively improves during hematopoietic stem cells to neutrophils differentiation

**DOI:** 10.64898/2026.09.03.749278

**Authors:** Allen Yesin, Kennedy Outlaw, Regina Sanchez-Flores, Aichatou Doucoure, Ruth Hüttenhain, Hawa Racine Thiam

## Abstract

Neutrophils’ ability to rapidly and efficiently migrate through narrow pores in tissues is essential for host defense and is proposed to depend on their multilobulated and deformable nucleus. When during neutrophil differentiation does optimal migration and its proposed nuclear determinants emerge, and whether they are intrinsic to progenitors, is unclear in part because of the scarcity of tractable models of human neutrophils and their progenitors. Here, we optimized a CD34+ hematopoietic stem cell (HSC) to neutrophil differentiation pipeline to generate millions of mature neutrophils (HSC-neutrophils) that recapitulate the surface markers, proteome, ROS production and NETosis of primary human blood-derived neutrophils, better than the widely used HL60-derived neutrophils. Comparative proteomics across differentiation showed that HSC-neutrophils become translationally repressed while acquiring immune functions and actin-related processes. Quantitative microscopy and proteomics showed that nuclear multilobulation occurs at the granulocyte progenitors-early neutrophils transition and is accompanied by drastic remodeling of nuclear envelope composition (increasing LBR, decreasing lamin A/C, B1/B2 and NUPs). Single nuclear envelope proteins only weakly correlate with nuclear multilobularity suggesting that an ensemble envelope state, rather than any one protein, sets nuclear shape. Using microfabricated devices with constrictions, we show that migration speed increases the most in early neutrophils; that the capacity to cross nucleus-deforming pores is continuously enhanced during differentiation; and that early neutrophils recover the best from such migration. We show that while mature neutrophils most effectively cross pores, they remain impaired. Our work resolves three features of neutrophils migration - speed, deformation through constrictions, and recovery from deformation - and maps when each emerges, opening the door to future mechanistic and engineering studies for modulating neutrophil migration in tissue-like microenvironments.

## Introduction

Neutrophils are the most abundant immune cells in our body and their rapid recruitment to tissues is critical for pathogen clearance during infection (1, 2). Neutrophils’ ability to optimally migrate through narrow pores, omnipresent in tissue, is proposed to maximize their recruitment to inflammation sites (3). This optimal migration requires three fundamental capabilities: rapid migration, effective deformation through constrictions smaller than the nucleus, and rapid recovery from such deformation. The multilobulated nucleus of mature neutrophils has been proposed to optimize rapid migration and deformation through narrow pores (4, 5). Indeed, while most cells have round nuclei, mature human neutrophils nuclei harbor 2-5 lobes which are proposed to provide less steric hindrance allowing rapid migration through narrow pores (3). An alternative model proposed that their low expression of lamin A/C, independently of multilobulation, allows neutrophil deformation through medium sized pores (6, 7). A major open question in the field is how nuclear envelope composition relates to nuclear shape and optimal neutrophil migration through narrow pores where nuclear deformability limits migration and rapid recovery is important for sustained migration (3). Answering this question is important to advance our understanding of neutrophil biology with direct disease relevance.

A range of human diseases, including acute infection, systemic stress, and cancer, coincides with abnormal recruitment of neutrophil progenitors to the blood stream and tissues (8, 9). Once in tissues, neutrophil progenitors can recapitulate the functions of primary neutrophils, including reactive oxygen species (ROS) production, phagocytosis and neutrophil extracellular trap release (NETosis) (8, 10). Yet, it remains unclear whether these neutrophil progenitors intrinsically possess mature neutrophil functions or acquire them upon recruitment to tissues. In particular, we do not know whether neutrophil progenitors migrate as effectively through narrow pores as mature neutrophils, nor how their nuclear shape and composition impact their migration in tissue-like environments. A fundamental question is therefore, when during neutrophil differentiation does optimal migration and its proposed nuclear determinants emerge, and whether these are intrinsic to progenitors or acquired only upon maturation.

Studying the migration behaviors of mature human neutrophils and progenitors has been limited by the scarcity of tractable in vitro models of these cells. Primary human neutrophils are most relevant, but they are short lived both in vivo and in vitro and accessing their progenitors requires the difficult and unscalable isolation of these cells from the bone marrow. HL60-derived neutrophils(dHL60 cells) have been extensively used to study neutrophil functions including migration and NETosis. For instance, studies in dHL60 cells revealed that neutrophils downregulate lamin A/C and B1 and LINC complex proteins (11–13) and this downregulation of lamin A/C allows them to optimally migrate through 3 µm pores (6). Their genetic tractability also helped to establish that the cellular events of NETosis are conserved and that PAD4 is important in human neutrophil NETosis (14, 15). However, dHL60 cells do not fully recapitulate the proteome, nuclear envelope composition or nuclear multilobulation of primary neutrophils (16–19), making them suboptimal for studying the emergence and impact of nuclear multilobulation on neutrophil migration.

More recently, stem cell-derived neutrophils differentiated from human induced pluripotent stem cells (iPSCs) or CD34+ hematopoietic stem cells (HSCs) have emerged as renewable, genetically tractable models that reproduce the maturation, antimicrobial activity, and granule biology of primary blood neutrophils (20–24). Since the differentiation proceeds through defined developmental stages, these cell models offer a unique opportunity to study the migration behaviors of human neutrophils and their progenitors and how these behaviors couple with nuclear remodeling.

Here, we adapted an HSC-neutrophil differentiation pipeline to generate large numbers of mature human neutrophils that, compared to dHL60 cells, better recapitulate the surface markers and whole proteome of primary neutrophils. Functionally, our HSC-neutrophils recapitulate the ROS production and NETosis ability of primary human neutrophils exposed to stimulants that mimic sterile inflammation and bacterial infection. Using comparative proteomics across different stages of HSC to neutrophil differentiation, we showed that molecular features of translation repression and immune responses emerge at the granulocyte progenitor-neutrophil transition whereas actin-related functions emerge late during neutrophil differentiation. We further showed that most remodeling of nuclear shape and envelope composition occurs at the granulocyte progenitor-early neutrophil transition, and that perinuclear lamin B receptor (LBR), lamin A/C, B1 and B2 correlate only weakly with nuclear shape at the single-cell level. Using microfabricated devices, we showed that rapid migration is optimized in early neutrophils while constriction-passage is continuously enhanced as neutrophils mature. Additionally, we show that migration through pores that require nuclear deformation, alters granulocyte progenitors’ migration, long term, while minimally impacting neutrophil migration. Together, our data untangles three fundamental aspects of neutrophil optimal migration: the ability to migrate rapidly; the ability to deform through constrictions smaller than the nucleus; the ability to rapidly recover from such deformation. By revealing when these behaviors emerge in parallel with proteome restructuring during differentiation, our work opens the doors to new strategies for independently tuning the three fundamental aspects of optimal migration of human neutrophils and their progenitors in tissue like environments as needed by the disease state.

## Results

### In vitro differentiation generates HSC-neutrophils that better recapitulate the proteome of primary human neutrophils

To generate human neutrophils at different stages of differentiation, we established a cytokine-guided in vitro neutrophil differentiation protocol from human bone marrow-derived CD34+ hematopoietic stem cells (HSCs). We adapted a protocol previously developed by Kuhikar et al. (25) by incorporating modifications such as the use of StemSpan™ SFEM II as the basal medium throughout the entire differentiation process; addition of small molecule agonists to sustain HSC stemness; and adjusted concentrations of G-CSF to optimize neutrophil yield and development (see Materials and Methods). First, CD34+ HSCs were expanded for seven days in StemSpan SFEM II media supplemented with stem cell expansion factors to preserve stemness, yielding a 15- to 20-fold expansion. Second, expanded CD34+ HSCs were advanced into a three-phase differentiation protocol (**Figure 1A**). During phase one, cells were exposed for four days to cytokines that support differentiation toward myeloid progenitors (see Materials and Methods). During phase two, cells were exposed for three days to cytokines that support granulocyte differentiation (see Materials and Methods). During phase three, cells were exposed for seven days to G-CSF to support final neutrophil differentiation and maturation. This workflow produced a cumulative 72.8 ± 4.8-fold cellular expansion between the onset and end of differentiation (**Figure 1B**), with a cellular viability of 76.6 ± 7.6% and 90.2 ± 2.9% at day 14 and 21, respectively, (**Figure S1**), allowing us to produce millions of cells with segmented, multilobed nuclei (**Figure 1A**), hereafter HSC-derived neutrophils (HSC-neutrophils) in 15 days.

**Figure 1:**
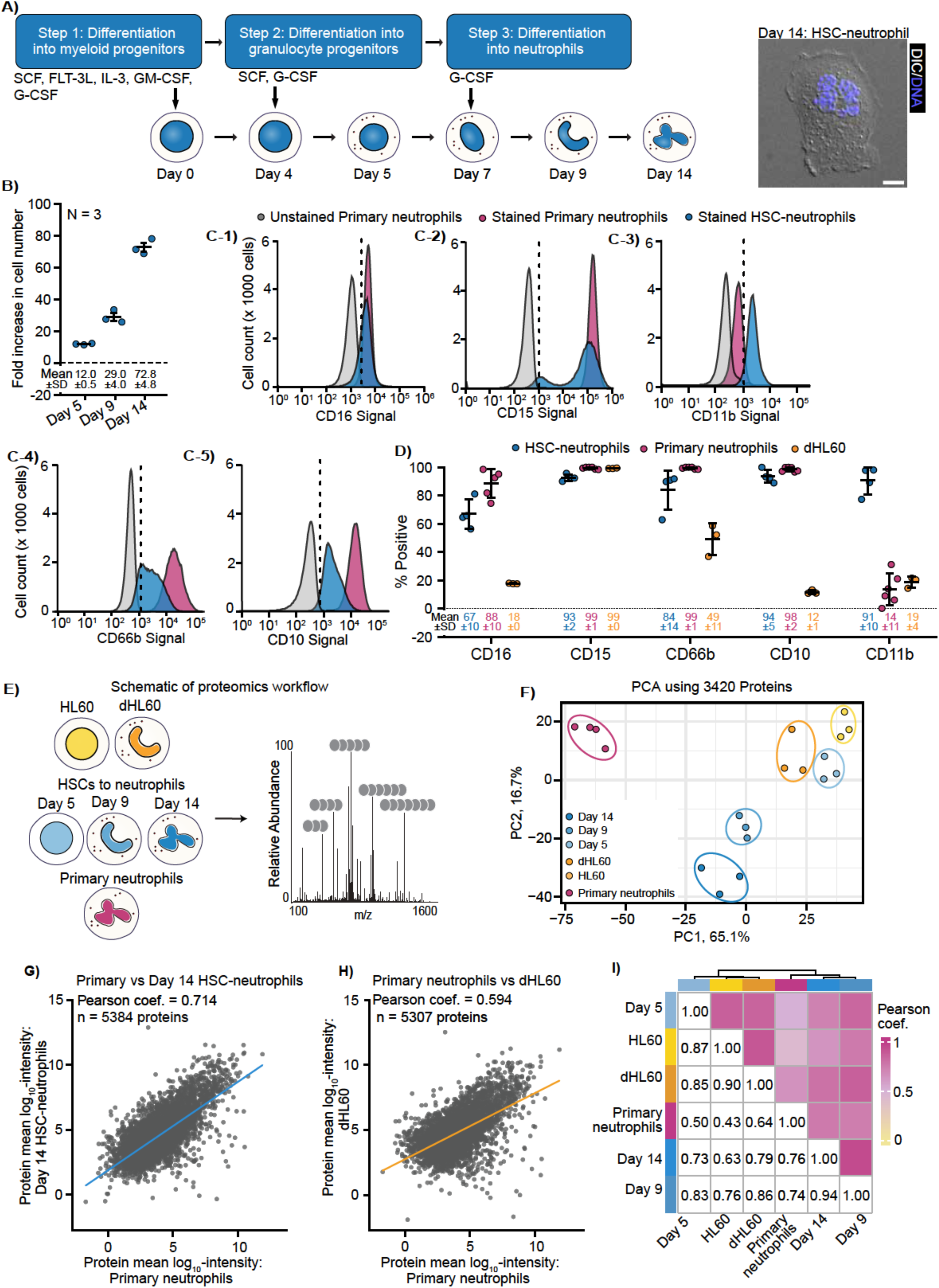
CD34+ hematopoietic stem cells can be differentiated into mature neutrophils that better recapitulate primary human neutrophils surface markers and proteome. **(A)** Schematic of the three-step CD34+ HSC to neutrophil differentiation protocol. Step 1 (Days 0 - 4): addition of SCF, FLT-3L, IL-3, GM-CSF and G-CSF for CD34+ HSCs into myeloid progenitor differentiation. Step 2 (Days 5 - 7): addition of SCF and G-CSF for myeloid to granulocyte progenitor differentiation. Step 3 (Days 9 - 14): addition of G-CSF for granulocyte progenitor to mature neutrophil differentiation. Right: representative image of a Day 14 HSC-neutrophil showing an overlay of DIC (grayscale) and DNA (blue) stained with SPY-650 DNA. **(B)** Fold increase in cell number at Day 5, Day 9 and Day 14 relative to number of cells at Day 0. N = 3 independent differentiations. **(C-1 - C-5)** Representative flow cytometry histograms of single-antibody stained CD16 **(C-1)**, CD15 **(C-2)**, CD11b **(C-3)**, CD66b **(C-4)** and CD10 **(C-5)** signal in unstained primary neutrophils (gray), stained primary neutrophils (red) and stained Day 14 HSC-neutrophils (blue). Dashed line indicates the positivity gate set on the unstained primary neutrophil control; positivity reflects gate crossing. Note that CD10 andCD66b peaks are left-shifted, suggesting their lower surface expression in HSC-neutrophils, compared to primary neutrophils. Histograms are representative of N ≥ 3 biological replicates. **(D)** Percentage of Day 14 HSC-neutrophils (blue), primary neutrophils (red) and dHL60 cells (orange) positive for CD16, CD15, CD11b, CD66b, CD10, gated on the unstained primary neutrophils control. N = 3 (dHL60), N = 4 (HSC-neutrophils) and N ≥ 5 (primary neutrophils), with n ≥ 10,000 events acquired per sample. **(E)** Schematic of proteomics workflow. Global proteomic composition of HL60, dHL60, HSC-neutrophils at Day 5, Day 9, and Day 14, and primary neutrophils were analyzed using mass spectrometry. **(F)** Principal component analysis (PCA) HL60, dHL60, HSC-neutrophils at Day 5, Day 9, and Day 14, and primary neutrophils. **(G)** Pearson correlation analysis of the mean log_10_ intensities between primary neutrophils and HSC-neutrophils (Day 14), n = 5384 proteins. **(H)** Pearson correlation analysis of the mean log_10_ intensities between primary neutrophils and dHL60 cells, n = 5307 proteins. **(I)** Pearson correlation analysis matrix of HL60, dHL60, HSC-neutrophils at Day 5, Day 9, and Day 14, and primary neutrophils showing replicate correlations averaged into a single box. Heat map to visualize Pearson coefficient from 0 < r < 1. **(B, D, F)** each point represents an independent biological replicate. Pearson coef. (for coefficient) shown on graphs for **(G)** and **(H)**. Bars in **(B)** and **(D)** indicate mean ± SD, and values shown below graph. (**E, F**). N = 3 (HSC-neutrophils, dHL60, HL60); N = 4 (primary human neutrophils). Scale bar in **(A)**: 5 μm.

We next assessed how closely HSC-neutrophils recapitulate the surface marker profile, proteome, and cellular functions of human peripheral blood-derived neutrophils (hereafter primary neutrophils). We also benchmarked HSC-neutrophils against dHL60 cells, the most commonly used human neutrophil-like cell line (26). We prioritized proteome-over transcriptome-based comparison with the reasoning that neutrophils are short-lived (27, 28); suggested to be transcriptionally repressed (29, 30); and functional characterization of HSC-neutrophils required mapping of proteins that mediate cellular functions.

First, we tested if HSC-neutrophils express common neutrophil surface markers to levels similar to what is detected in primary human neutrophils. We focused on five surface markers commonly used to characterize neutrophil identity and maturation: CD16, an Fcγ receptor expressed by neutrophils, macrophages and NK cells (31); CD15, a carbohydrate present in neutrophils and myeloid cells (32); CD11b, an integrin present in myeloid cells including neutrophils (33); CD66b, a GPI-anchored glycoprotein specific to neutrophils and eosinophils (34, 35); and CD10, a metalloprotease specific to mature neutrophils (36, 37). We used flow cytometry to measure the expression levels of these surface markers in HSC-neutrophils, primary neutrophils and dHL60 cells. Analysis of the flow cytometry profiles indicated that HSC-neutrophils express CD16, CD15, CD66b and CD10 at levels comparable to primary neutrophils (**Figure 1C, D**). Specifically, ∼67%, ∼93%, ∼84% and 94% of HSC-neutrophils highly express CD16, CD15, CD66b, and CD10, respectively (**Figure 1D**). This is comparable to the ∼88%, ∼99%, ∼99% and ∼98% of primary neutrophils that express CD16, CD15, CD66b, and CD10, respectively. Notably, ∼91% of HSC-neutrophils express CD11b while only ∼14% of primary neutrophils express this surface marker (**Figure 1D**), which is consistent with previous reports that peripheral blood-derived neutrophils minimally expose CD11b to their surface (38–40). Importantly, dHL60 cells lowly express CD16 (∼18%), CD66b (∼49%) and CD10 (∼12%) (**Figure 1D, Figure S2**), indicating that they poorly mirror the surface molecule profile of primary neutrophils. These data indicate that our differentiation protocol yields mature neutrophils with surface markers similar to that of primary human neutrophils.

Second, to assess if HSC-neutrophils better recapitulate primary neutrophils’ proteome, compared to dHL60 cells, we performed global proteome profiling using quantitative proteomics (see Materials and Methods) comparing primary neutrophils, dHL60 cells and HSC-neutrophils across differentiation (**Figure 1E**). We focused on Day 5 (24hr after induction of granulocyte progenitors), Day 9 (48hr after induction of neutrophils) and Day 14 (HSC-neutrophils) cells. We quantified more than 75,000 peptides and over 5,700 unique protein groups per cell type across all replicates with primary and HSC-neutrophils having the lowest number of unique protein groups (5,728 and 6,960 proteins, respectively; **Figure S3 A-C**). Principal component (PC) analysis revealed tight clustering of replicates and a progressive trajectory across differentiation: Day 5, 9, and 14 cells separated along PC1 with decreasing interstage distances, consistent with reproducible, stepwise differentiation (**Figure 1F**). Day 5 cells clustered with HL60 cells, whereas Day 9 and 14 cells remained distant from dHL60 cells. Notably, HSC-neutrophils (Day 14) and primary neutrophils were closest along PC1 which captured the majority (65%) of the variance, indicating that Day 14 HSC-neutrophils better recapitulate primary neutrophils’ proteome than dHL60 cells. This was confirmed by quantitative correlation analysis, hierarchical clustering and comparison of the protein expression levels of six markers commonly used to characterize neutrophil identity and maturation. The mean abundance of proteins shared between primary and Day 14 HSC-neutrophils (n = 5384 proteins; Pearson coef. = 0.714, **Figure 1G, I**; Spearman coef. = 0.660, **Figure S3D, F**) showed a higher correlation compared to proteins shared between primary neutrophils and dHL60 cells (n = 5307 proteins, Pearson coef. = 0.594, **Figure 1H, I**; Spearman coef. = 0.538, **Figure S3E, F**). Plus, hierarchical clustering of the full correlation matrix showed that Day 5, Day 9 and Day 14 cells cluster the closest to HL60, Day 14 HSC-neutrophils and primary neutrophils, respectively (**Figure 1I and S3F**). Analysis of the abundance (mean log_2_ intensity) of progenitor markers (c-kit and FUT4 (41, 42)), early neutrophil markers (CD66b and CD11b) and late stage, mature neutrophil markers (CD16 and CD10) showed that progenitor markers are abundant in Day 5 cells and not detected in Day 14 cells (**Figure S3G-1,2**); neutrophil markers increase drastically between Day 5 and Day 9 cells (**Figure S3G-3,4**); and mature neutrophil markers are not or minimally detected in Day 5 and Day 9 cells, and drastically increase in Day 14 cells and primary neutrophils (**Figure S3G-5,6**). These data indicate that Day 5, 9 and 14 cells are granulocyte progenitors, early and mature neutrophils, respectively and that Day 14 HSC-neutrophils better recapitulate primary neutrophil proteome, compared to dHL60 cells.

### HSC-neutrophils functionally recapitulate primary human neutrophils ROS production and NETosis behavior

We next sought to determine whether HSC-neutrophils functionally recapitulate primary human neutrophils by assessing their ability to undergo respiratory burst and neutrophil extracellular traps release (NETosis) upon stimulation, two critical functions of neutrophils.

To test if HSC-neutrophils undergo respiratory burst we compared their ability to produce reactive oxygen species (ROS) upon stimulation with Phorbol 12-myristate 13-acetate (PMA), a well-established neutrophil activator (43, 44). Cells were stained with DCFDA (2’,7’-dichlorofluorescin diacetate; a cell permeant probe that once oxidized produces the fluorescent compound DCF (45)) then stimulated with PMA (20 nM) or vehicle control (DMSO). Unstained and DCFDA-stained cells were analyzed by fluorescence endpoint assay using a plate reader, 30 minutes after the onset of PMA stimulation (see Materials and Methods). Observation via widefield microscopy (**Figure 2A, Figure S4**) and quantification (**Figure 2B**) of the DCF fluorescent signal showed a significant, ∼1.77-fold, increase in HSC-neutrophils upon PMA treatment. This increase is within range of the non-significant ∼1.69-fold increase in DCF signal measured in primary neutrophils upon PMA treatment. This data indicate that HSC-neutrophils can produce ROS upon PMA stimulation.

**Figure 2:**
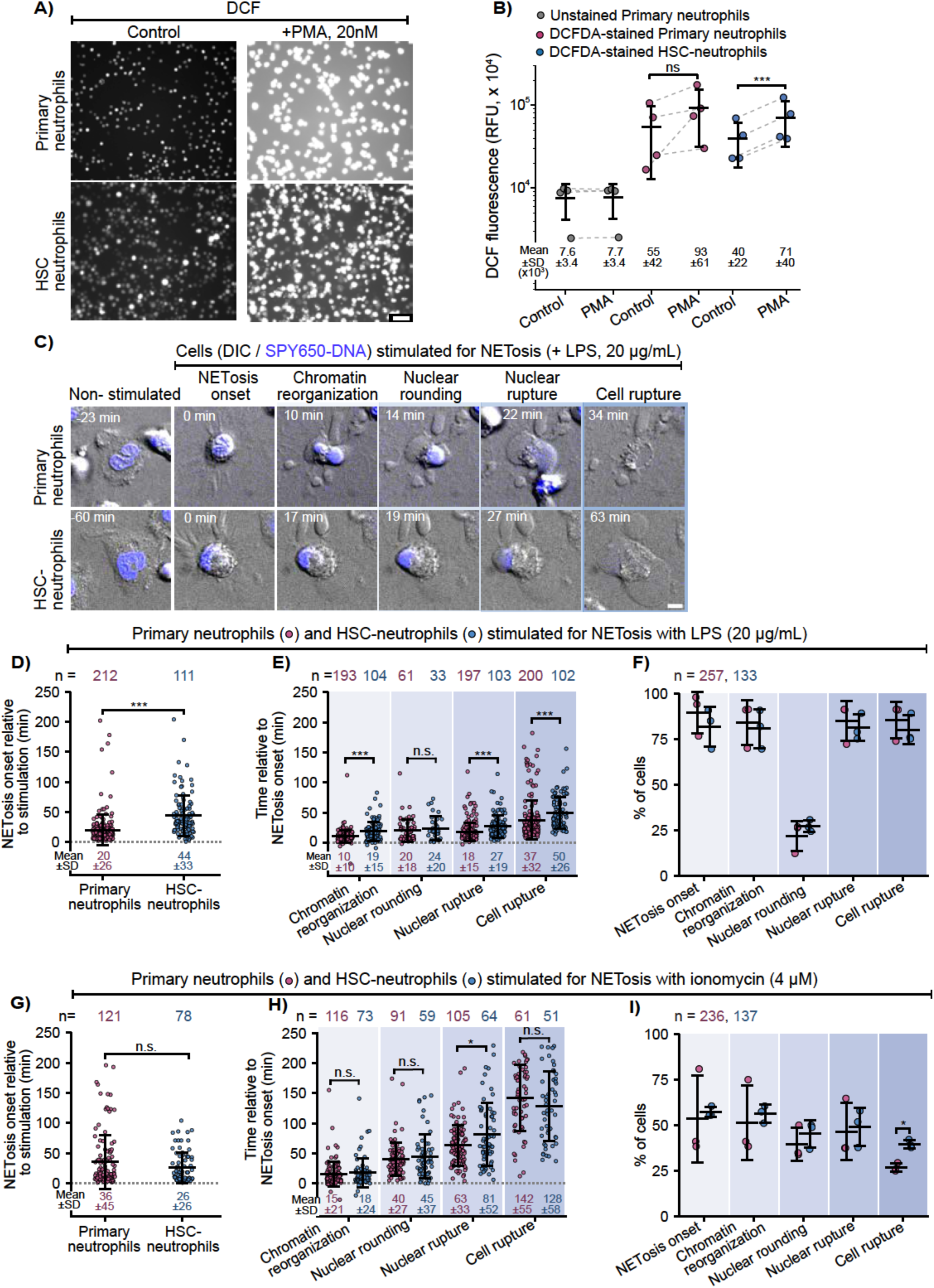
HSC-neutrophils recapitulate primary neutrophils ROS production and NETosis progression. **(A)** Representative 20x widefield microscopy images of DCF fluorescence in DCFDA-loaded primary neutrophils (top) and HSC-neutrophils (bottom) treated with DMSO (Control) or PMA (20 nM). **(B)** Quantification of total DCF fluorescence (RFU × 10⁴) via fluorescence endpoint plate reader assay in unstained (gray) and stained (red) primary neutrophils and stained HSC-neutrophils (blue), control or PMA-treated. Gray dashed lines to connect biological replicates. N = 3 independent experiments. **(C)** Representative montages of primary (top) and HSC-neutrophils (bottom) stimulated for NETosis with LPS (20 μg/mL) and quantified in (D-F). Images show overlays of DIC (grayscale) and SPY650-DNA (blue) at the indicated stages of NETosis: non-stimulated, NETosis onset, chromatin reorganization, nuclear rounding, nuclear rupture and cell rupture. **(D)** Quantification of timing of NETosis onset relative to stimulation with LPS (20 μg/mL) in primary (red) and HSC-neutrophils (blue). **(E)** Quantification of timing of stages represented in (C) relative to NETosis onset, in cells stimulated with LPS. **(F)** Percentage of LPS-stimulated cells at different stages of NETosis. **(G–I)** Quantification of timing of NETosis onset relative to ionomycin stimulation (4 µM, **G**); timing of NETosis stages represented in (C) relative to NETosis onset (**H**); percentage of ionomycin stimulated cells at different stages of NETosis (**I**). Brackets denote the presence of ionomycin in **(D - F)** and of LPS in **(C)** and **(G - I)**. **(D - I)**: n represents number of cells. Bars in **(B-I)** indicate mean ± SD, and values shown below graph for (**B, D, E, G, H**). Statistical test in **(B)**: paired t-test; **(D, E, G, H)**: Mann Whitney; in **(F, I)** Fischer test; n.s.: not significant; *: p-value < 0.05; ***: p-value < 0.001; ****: p-value < 0.0001. (**F, I**) all statistical tests were not significant unless indicated. Scale bars: **(A)** 40 μm; **(C)** 5 μm

To test if HSC-neutrophils undergo NETosis as efficiently as primary neutrophils, we compared their ability to initiate and execute the cellular events leading to NETosis following stimulation with two well established NETosis stimulants, lipopolysaccharide (LPS) from the bacteria *Klebsiella pneumoniae* (20 ug/mL) and ionomycin (4 µM) (15, 44). Cells were stained with SPY650-DNA to visualize the nucleus, then imaged on a spinning disk confocal microscope for 4 hours, with NETosis stimulants added five minutes after imaging onset. Observation (**Figure 2C, Figure S5, Movie S1-S2**) and analysis of time-lapse movies (**Figure 2D-I**) showed that both HSC- and primary neutrophils progressed through NETosis following the same sequence of cellular events, consistent with our previous work (15, 44, 46).

LPS-stimulated HSC- and primary neutrophils initiated NETosis by retracting their membrane protrusions before shedding microvesicles, reorganizing their chromatin, and then rupturing their nuclear and plasma membranes to allow NET release (**Figure 2C, Movie S1**). Quantification of the time of NETosis onset relative to stimulation (**Figure 2D**) as well as the times between NETosis onset and chromatin reorganization, nuclear and plasma membrane rupture showed that HSC-neutrophils initiate and progress through LPS-induced NETosis significantly slower than primary neutrophils (**Figure 2E**). Importantly, quantification of the percentage of cells that progress through each cellular event showed no significant difference between the two cell types (**Figure 2F**) indicating that the slower NETosis kinetics of HSC-neutrophils does not significantly impact their NETosis probability.

Ionomycin-stimulated HSC- and primary neutrophils initiated NETosis by first shedding microvesicles from the plasma membrane, then reorganizing their chromatin, rounding and rupturing their nuclei, and finally rupturing their plasma membrane (**Figure S5, Movie S2**). Quantification of the time of NETosis onset relative to stimulation (**Figure 2G**), as well as the times between NETosis onset and chromatin reorganization, nuclear rounding and cell rupture (**Figure 2H**) showed that HSC-neutrophils initiate and progress through ionomycin-induced NETosis with similar kinetics to primary neutrophils. Plus, quantification of the percentage of cells that progress through each NETosis stage showed that a similar percentage of HSC-neutrophils initiate and progress through NETosis within 4 hours, compared to primary neutrophils (**Figure 2I**). Importantly, while HSC-neutrophils rupture their nucleus later compared to primary neutrophils (**Figure 2H**), they rupture their cell membrane and release NETs at a higher percentage (**Figure 2I**), indicating an overall higher NETosis efficiency.

This data indicates that HSC-neutrophils recapitulate two critical functions of primary human neutrophils, respiratory bursts and the release of neutrophils extracellular traps following two distinct stimulations.

Our data so far show that HSC to neutrophils differentiation produces cells at distinct stages of neutrophil differentiation and mature neutrophils that proteomically and functionally recapitulate primary human neutrophils. This gives us access to mature neutrophils and neutrophil progenitors to interrogate when neutrophil behaviors including optimal migration emerge during neutrophil differentiation.

### HSC-neutrophil differentiation is accompanied by translation repression and acquisition of immune and actin-regulated functions

To start probing when neutrophil behaviors, including optimal migration, emerge during neutrophil differentiation, we leveraged our proteomics dataset to assess proteome changes across granulocyte progenitors (Day 5), early neutrophils (Day 9) and mature HSC-neutrophils (Day 14).

To assess proteome changes during neutrophil differentiation, we performed pairwise comparisons of the distinct differentiation stages (Day 9 vs Day 5; Day 14 vs Day 5; and Day 14 vs Day 9). We then filtered the results for proteins that are significantly differentially expressed in at least one comparison (adj-p-value<0.05) and are detected across all differentiation stages and replicates, which resulted in 3877 differentially expressed proteins (DEP). Hierarchical clustering of their z-score across cell types resolved four clusters, which were subjected to gene ontology (GO) enrichment analysis of biological processes (**Figure 3A**). Cluster 1 (2,236 proteins; ∼58% of DEP), containing proteins highest in progenitors and progressively decreasing during differentiation, was enriched for RNA biogenesis and processing, translation and chromosome organization. Cluster 2 (665 proteins; ∼17% of DEP), containing proteins that were selectively reduced in early neutrophils, was enriched for ubiquitin-dependent protein regulation (including degradation), vesicle organization and trafficking, suggesting these processes are downregulated at the granulocyte-early neutrophil transition before partially recovering in mature neutrophils. Cluster 3 (230 proteins; 6% of DEP), representing proteins that increase in early neutrophils and are sustained in mature neutrophils, was enriched for phagocytosis, immune response and locomotion. Finally, cluster 4 (746 proteins, ∼19% of DEP), representing proteins that progressively increase during differentiation, was enriched for actin-based processes, intracellular signaling and myeloid cell activation. Together the data indicate that the translational activity of HSCs decreases while immune functions are acquired as they differentiate into neutrophils. Further, the gradual increase of actin-associated proteins suggests that actin-regulated processes, such as migration, are progressively upregulated during differentiation.

**Figure 3:**
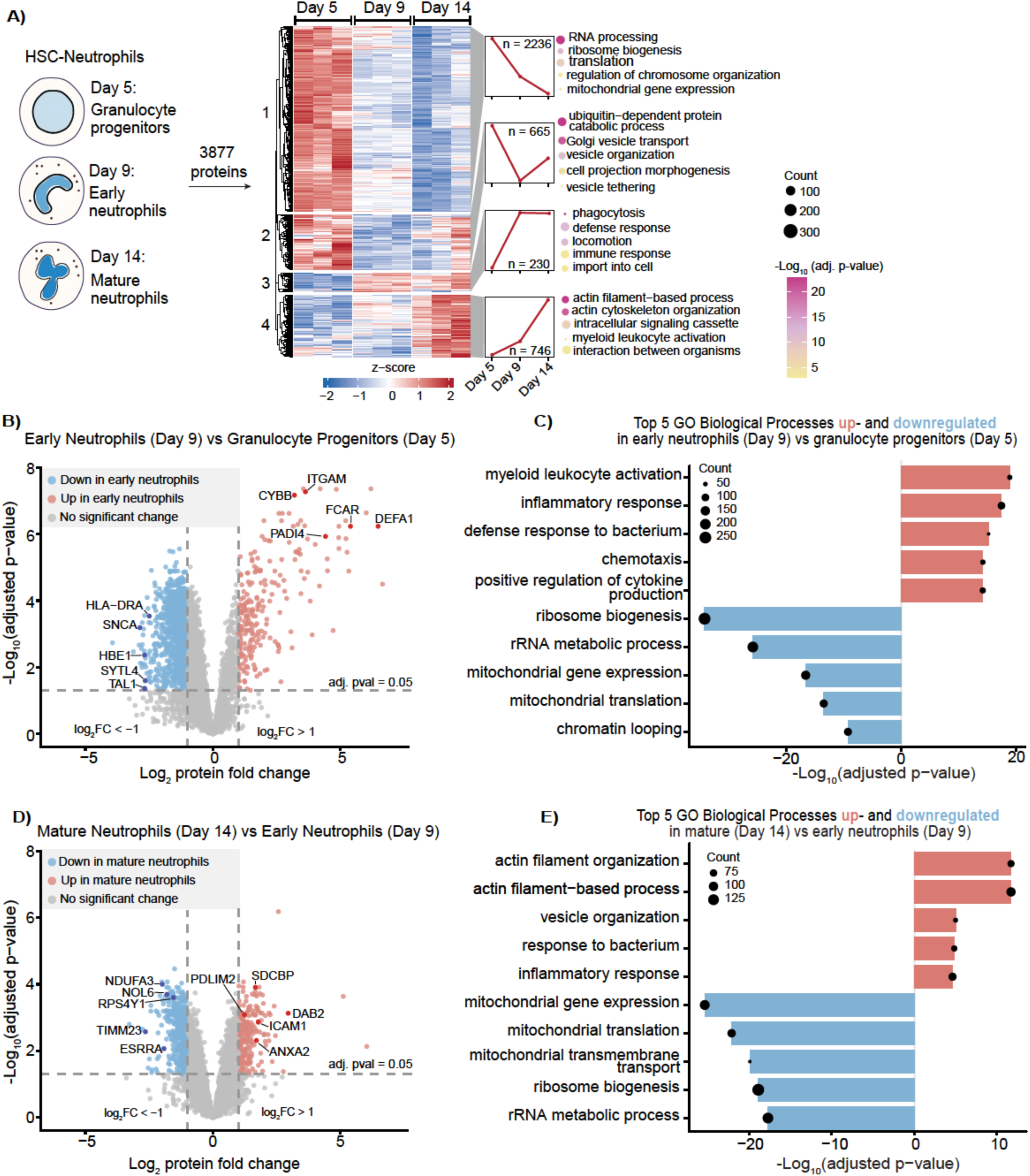
HSC-neutrophils become translationally repressed and gain immune and actin regulated functions. **(A)** HSC-neutrophils at Day 5 (Granulocyte progenitors), Day 9 (Early neutrophils), and Day 14 (Mature neutrophils) were analyzed via mass spectrometry (Fig 1E). Proteins significantly changing (p.adjusted < .05) in at least on pairwise comparison, (Day 5 vs Day 9, Day 9 vs Day 14 or Day 5 vs Day 14 cells), where filtered for presence in each sample (n = 3,877 proteins) and depicted in a heat map colored by z-score (z-score ± 2) and separated into four clusters by hierarchical clustering. Z-score trajectories are shown next to each cluster along with the number of proteins in that cluster (Cluster 1; n = 2,236 proteins), (Cluster 2; n = 665 proteins), (Cluster 3; n = 230 proteins), (Cluster 4; n = 746 proteins). Gene Ontology (GO) enrichment is shown next to each cluster representing the top enriched biological processes colored by significance (-log_10_ adjusted p-value ∈ [5, 20]) and gene count. **(B)** Volcano plot showing differential protein abundance for Early neutrophils vs Granulocyte progenitors (|log₂ FC|≥ 1, adjusted p-value ≤ 0.05). 5 representative proteins from both up regulated (red), and down regulated, (blue) are shown to illustrate GO pathways shown in C. **(C)** GO enrichment showing the top 5 most enriched biological processes within the upregulated (red) and downregulated (blue) proteins in early neutrophils. **(D)** Volcano plot showing differential protein abundance for Early vs Mature neutrophils (|log₂ FC|≥ 1, adjusted p-value ≤ 0.05). 5 representative proteins from both up regulated (red), and down regulated(blue) are shown to illustrate GO pathways shown in E. **(E)** GO enrichment showing the top 5 most enriched biological processes within the upregulated (red) and downregulated (blue) proteins in mature neutrophils.

To pinpoint at which cellular transition these proteome changes occur, we analyzed the individual pairwise comparisons between granulocyte progenitors (Day 5) and early neutrophils (Day 9) (**Figure 3B, C**) and between early (Day 9) and mature neutrophils (Day 14) (**Figure 3D, E**). At the granulocyte progenitor-early neutrophil transition, 1,291 and 235 proteins were down- and upregulated, respectively (|log_2_ FC| ≥ 1; adj-p-value ≤ 0.05; **Figure 3B**). GO analysis for biological processes showed that downregulated proteins were enriched for RNA biogenesis and processing, mitochondrial gene expression and chromatin looping, while the upregulated proteins were enriched for immune processes including myeloid cell activation, defense response to bacterium, inflammatory response, cytokine production and chemotaxis (**Figure 3C**). Thus, the transition between granulocyte progenitors and early neutrophils is most characterized by depletion of translation machinery and the emergence of immune processes.

At the early-mature neutrophil transition, 469 and 228 proteins were down- and upregulated, respectively (|log_2_ FC| ≥ 1; adj-p-value ≤ 0.05; **Figure 3D**). Downregulated proteins participated in RNA processing but also in mitochondria gene expression and membrane transport, indicating that both translation/transcription dynamics and mitochondrial functions are tuned in neutrophils (**Figure 3E**). Critically, upregulated proteins were enriched for actin filament organization and actin-regulated processes (**Figure 3E**) as well as vesicle organization and immune response, identifying the early to mature transition as the stage in which most of the actin-regulated processes are upregulated.

Analysis of the pairwise comparison between granulocyte progenitors (Day 5) and mature neutrophils (Day 14) confirmed that the most downregulated proteins (n = 1,976) relate to translation and chromatin looping while the most upregulated proteins (n = 495) relate to actin organization, actin-based processes and immune response (**Figure S6**).

Together, this data shows that neutrophil differentiation is accompanied by progressive downregulation of the translation machinery and upregulation of immune response and actin-based processes. Our finding that chemotaxis and locomotion are upregulated at the granulocyte progenitors to early neutrophil transition and that actin-based processes are progressively upregulated during the early to mature neutrophil transition suggests that cell migration machinery is in place by the early neutrophil stage and continues to be built through differentiation.

### Nuclear multilobulation is initiated in early neutrophils and only weakly correlates with LBR, lamin A/C, lamin B1 and lamin B2 expression in neutrophils

Neutrophils’ ability to rapidly deform through pores smaller than the nucleus, a critical feature of optimal migration, has been attributed to the distinct envelope composition and multilobulation of the neutrophil nucleus (3, 5, 6). We reasoned that determining when nuclear envelope composition and multilobulation change during neutrophil differentiation will allow us to generate hypotheses on whether optimal migration is established in neutrophil progenitors or is a feature of mature neutrophils.

We first mined our proteomics dataset for nuclear envelope proteins, as annotated by the Human Protein Atlas (n = 285 proteins, **Supplementary Table 1**, (47)). Of the 193 nuclear envelope proteins detected in our dataset, 119 were significantly differentially expressed based on pairwise comparison in Figure 3A (adj-p-value ≤ 0.05). Hierarchical clustering of their z-score across cell types resolved three main clusters (**Figure 4A**). Cluster 1 (n = 23 proteins; including LBR, CHMP2A, and TOR1AIP1) was progressively upregulated in early and mature neutrophils relative to granulocyte progenitors. Cluster 2 (n = 18 proteins, including TMPO, LEMD2/3 and RAN), was selectively downregulated in early neutrophils. Cluster 3, the largest, (n = 78 proteins; including LMNA, LMNB1/B2 and NUP155) was predominantly continuously downregulated in early and mature neutrophils relative to granulocyte progenitors. Mapping the significantly changing nuclear envelope proteins into the STRING interaction network, we found that proteins within the same cluster tend to share interaction networks (**Figure S7**). Notably, downregulated proteins in cluster 3 are associated with the nuclear pore complex, nuclear transport and nuclear lamina. Upregulated proteins in cluster 1 are associated with the ESCRT complex. This indicates that different biological processes, occurring at the nuclear envelope, are dynamically remodeled during neutrophil differentiation.

**Figure 4:**
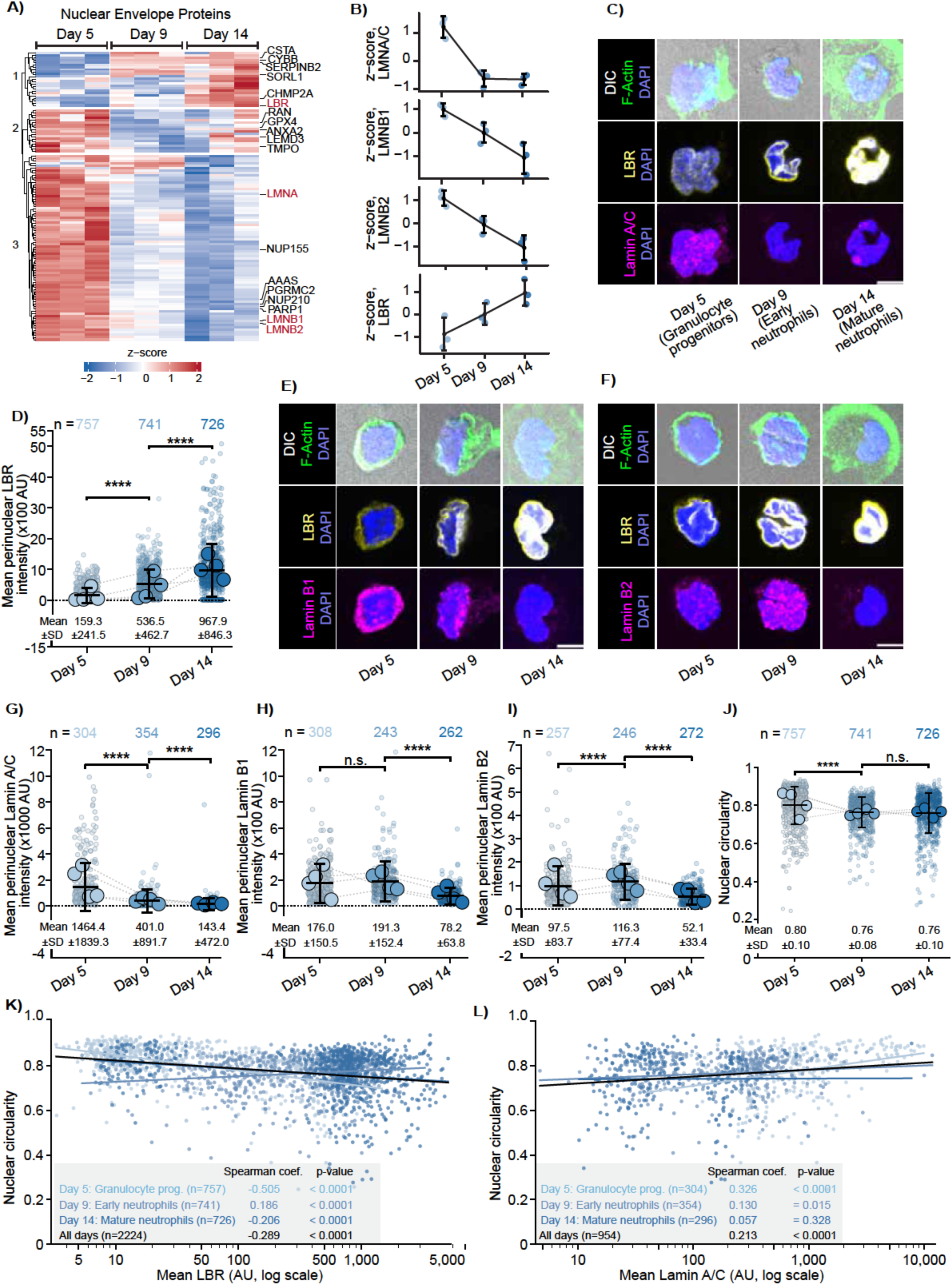
Nuclear shape and composition change is an early and progressive process during neutrophil differentiation. **(A)** Human Protein Atlas was used to curate a list of proteins to search the data set. The resulting heatmap of nuclear envelope proteins present in Day 5, Day 9, and Day 14 cells from Figure 3A was colored by z-score and separated into three clusters using hierarchical clustering. **(B)** Z-score trajectories of LMNA/C, LMNB1, LMNB2, and LBR in Day 5, 9 and 14 cells. **(C)** Representative images of Day 5, Day 9 and Day 14 cells showing (Top) DIC overlaid with F actin (Phalloidin, green) and DNA (DAPI, blue); (Middle) LBR (yellow) with DAPI; and (Bottom) lamin A/C (magenta) with DAPI. **(D)** Mean perinuclear LBR intensity (×100 AU) per nucleus in Day 5, Day 9 and Day 14 cells. **(E -F)** Representative images of Day 5, Day 9 and Day 14 cells showing (Top) DIC overlaid with F-actin (Phalloidin, green) and DNA (DAPI, blue); (Middle) LBR (yellow) with DAPI; and (Bottom) lamin B1 (magenta, **E**) or lamin B2 (magenta, **F**) with DAPI. **(G–I)** Mean perinuclear lamin A/C (×1000 AU; **G**), lamin B1 (×100 AU; **H**) and lamin B2 (×100 AU; **I**) intensity per nucleus in Day 5, Day 9 and Day 14 cells. **(J)** Nuclear circularity in Day 5, Day 9 and Day 14 cells. **(K, L)** Single-cell correlation between nuclear circularity and mean perinuclear LBR (**K**) or lamin A/C (**L**) intensities x-axis on a log scale. Lines indicate best fit for each day (color codes in blue) and for all days pooled (black); Spearman coefficients and associated p-values are reported in the inset table. Fluorescence intensities in **(D)** and **(G - I)** were measured as the mean pixel intensity per nucleus within a nuclear mask defined by the DAPI signal and are expressed in arbitrary units (AU). Data in **(D)** and **(G–J)** are shown as Superplots, where each small dot represents a cell and each large dot represents the mean of a biological replicate; N = 4 independent experiments/differentiations; n = total number of cells. Bars indicate mean ± SD, values shown below the graphs. Statistical test: Mann-Whitney on 2 compared groups using pooled single-cell data; n.s.: not significant; *: p-value < 0.05; ***: p value < 0.001; ****: p-value < 0.0001. Scale bars: 5 μm.

To map how well-established nuclear envelope proteins are tuned during neutrophil differentiation, we examined the z-score trajectories of lamin A/C, lamin B1/B2, and LBR (**Figure 4B**). This showed that while lamin A/C decreased abruptly in early neutrophils and remained stable as a low abundance protein, indicating downregulation at the onset of differentiation, lamin B1/B2 and LBR changed gradually (decreasing and increasing, respectively) throughout neutrophil differentiation. Thus, nuclear envelope proteins are dynamically regulated both at the onset of and throughout neutrophil differentiation.

To determine if these trends in nuclear envelope-associated protein abundance by bulk proteomics translate into changes in nuclear envelope composition and nuclear multilobulation at single cell level, we used quantitative microscopy to measure the perinuclear distribution of a select group of proteins (LBR, lamin A/C, B1 and B2) in relation to nuclear shape in cells at different stages of neutrophil differentiation. Granulocyte progenitors (Day 5), early (Day 9) and mature (Day 14) neutrophils were fixed and stained for DAPI and phalloidin to visualize the nucleus and actin cytoskeleton, respectively, and immunostained for LBR, lamin A/C, B1 or B2. Cells were imaged using a spinning disk confocal microscope. Z-series were acquired to obtain 3D distributions of our protein of interest. Observation and analysis of the mean fluorescence intensity of our proteins of interest at the nuclear periphery, as denoted by the DAPI signal (see Materials and Methods), showed that perinuclear LBR and lamin A/C gradually increased and decreased, respectively, as neutrophil differentiate (**Figure 4C, D, G, Figure S8A**). Comparison of the fold change in LBR revealed a ∼3-fold increase in perinuclear LBR at the granulocyte-early neutrophil transition, followed by a ∼1.8-fold increase at the early-mature neutrophil transition (**Figure 4D**), indicating that perinuclear LBR increases the most at the onset and not at the mature stage of neutrophil differentiation. Similar analysis of perinuclear lamin A/C showed a ∼3.6-fold decrease at the granulocyte-early neutrophil transition, followed by a ∼2.6-fold decrease at the early-mature neutrophil transition (**Figure 4G**), suggesting a progressive decrease in perinuclear lamin A/C throughout neutrophil differentiation.

Observation (**Figure 4E, F, Figure S8B, C**) and analysis (**Figure 4H, I**) of perinuclear lamin B1 and B2 revealed that these proteins most drastically change at the early-mature neutrophils transition. We measured a ∼1.08- and ∼1.2-fold increase in lamin B1 and B2, respectively, between granulocyte progenitors and early neutrophils followed by a ∼2.4- and ∼2.2-fold decrease between early and mature neutrophils. This data deviates from our proteomics analysis that indicated progressive decrease in lamin B1 and B2 throughout neutrophil differentiation (**Figure 4B**). To resolve this, we observed the whole cell distribution of lamin B1 and B2 and found a non-perinuclear pool of these proteins in granulocyte progenitors that is not detected in neutrophils (**Figure 4E, F, Figure S8B, C**). Together, these data indicate that the composition of the nuclear envelope, not only the expression of nuclear envelope proteins, is dynamically tuned as neutrophils differentiate and mature. These results further show that perinuclear LBR and lamin A/C change the most at the early stages of neutrophil differentiation while perinuclear lamin B1 and B2 decrease the most between early and mature neutrophils.

We next set out to determine if this change in nuclear envelope composition relates to nuclear multilobulation. To better capture the complexity of the nuclear periphery, we used LBR signal, rather than the DAPI signal, to generate a mask of the nuclear periphery and calculate nuclear circularity (see Materials and Methods). Comparison of cells across differentiation showed that nuclear circularity significantly decreased at the granulocyte progenitors-early neutrophils transition and does not significantly change between early and mature neutrophils (**Figure 4J**). This indicates that nuclear multilobulation is established early during neutrophil differentiation, coincident with the increase in perinuclear LBR and decrease in lamin A/C.

To determine if perinuclear LBR or lamin A/C correlates with nuclear multilobulation, at single cell resolution, we performed correlation analysis. Spearman correlation across granulocyte progenitors, early and mature neutrophils showed that perinuclear LBR intensity weakly but significantly negatively correlates with nuclear circularity (Spearman coef (All days) = -0.289; p-value < 0.0001; **Figure 4K**). The same analysis for lamin A/C showed that perinuclear lamin A/C weakly but significantly positively correlates with nuclear circularity (Spearman coef (All days) = 0.213; p-value < 0.0001; **Figure 4L**). Comparison of the Spearman correlation coefficients across differentiation stages revealed stages specific trends. The correlations between perinuclear LBR, lamin A/C and nuclear circularity are the strongest in granulocyte progenitors (Spearman coef = -0.505 for LBR and = 0.326 for lamin A/C in granulocytes). In early neutrophils that have the lowest nuclear circularity at the population level, perinuclear LBR does not negatively correlate with nuclear circularity (Spearman coef = 0.186; **Figure 4K**) and perinuclear lamin A/C weakly correlates with nuclear circularity (Spearman coef = 0.130; p-value = 0.015; **Figure 4L**). In mature neutrophils, perinuclear lamin A/C does not significantly correlate with nuclear circularity (Spearman coef. = 0.057; p-value = 0.328; **Figure 4L**). Correlation analysis between nuclear circularity and perinuclear lamin B1 and B2 showed that these proteins significantly positively correlate with nuclear circularity only in mature neutrophils (**Figure S8D, E**).

Together, our data indicate that both nuclear multilobulation and nuclear envelope composition are dynamically tuned during neutrophil differentiation. However, our analysis could not identify a strong correlation between nuclear multilobulation and a single nuclear envelope protein, at single cell level. Rather, our data identified specific trends for each differentiation stage, raising the possibility that nuclear envelope state, defined by an ensemble of proteins rather than a given protein, regulates nuclear multilobularity in neutrophils.

Our data so far indicate that granulocyte progenitors to early neutrophil differentiation is accompanied by downregulation of the translation machinery; upregulation of immune cell functions; acquisition of nuclear multilobulation and drastic remodeling of nuclear envelope composition, while the early to mature neutrophil transition is accompanied by increase in actin-based process and less drastic changes in nuclear shape and envelope composition. The decrease in nuclear circularity and in the expression level of many major nuclear structural proteins at the granulocyte progenitors-early neutrophil transition suggests an increase in nuclear deformability. We thus set out to test if migration that requires nuclear deformation is optimized in early neutrophils while rapid migration, powered by actin-based processes, is optimized in mature neutrophils.

### High deformability and optimal migration emerge early during neutrophil differentiation

To evaluate optimal migration (the ability to migrate rapidly, deform through constrictions smaller than the nucleus, and rapidly recover from such deformation) and determine when it emerges during neutrophil differentiation, we measured the migration behavior of granulocyte progenitors, early and mature neutrophils in two sets of microfabricated devices (**Figure 5**). Our first device consists of 7 µm wide and 5 µm tall channels that impose cellular confinement and allow the assessment of rapid migration, independent of nuclear deformation (**Figure 5A-top**). Our second device consists of 7 µm x 5 µm channels with 2 µm x 5 µm constrictions that impose nuclear confinement and allow the assessment of cellular deformation through constriction smaller than the nucleus and ability to rapidly recover from it (**Figure 5A-bottom**). Cells were stained with SPY650-DNA to visualize the nucleus, loaded in the inlet sections of the devices, left for 2-3 hours to spontaneously enter the channel section of the device then image for 8 hours with a spinning disk confocal microscope (see Materials and Methods). From the timelapse movies, we measured the cells’ instantaneous speed in channels; before, inside and after the constrictions; and the percentage of cells that pass constrictions (see Materials and Methods). Observation (**Figure 5B, Movie S3-S4**) and analysis of the distribution of cells’ average instantaneous speed in the 7 µm channels (**Figure 5C**) revealed that cells gradually increased their speed as they differentiate. Comparison of the fold change in average instantaneous speed showed a ∼1.6- and ∼1.1-fold increase in cell speed at the granulocyte progenitor-early neutrophil and the early-mature neutrophil transition, respectively. This indicates that cell speed increases the most at the granulocyte progenitors-early neutrophil transition, revealing that rapid migration emerges early during neutrophil differentiation.

**Figure 5:**
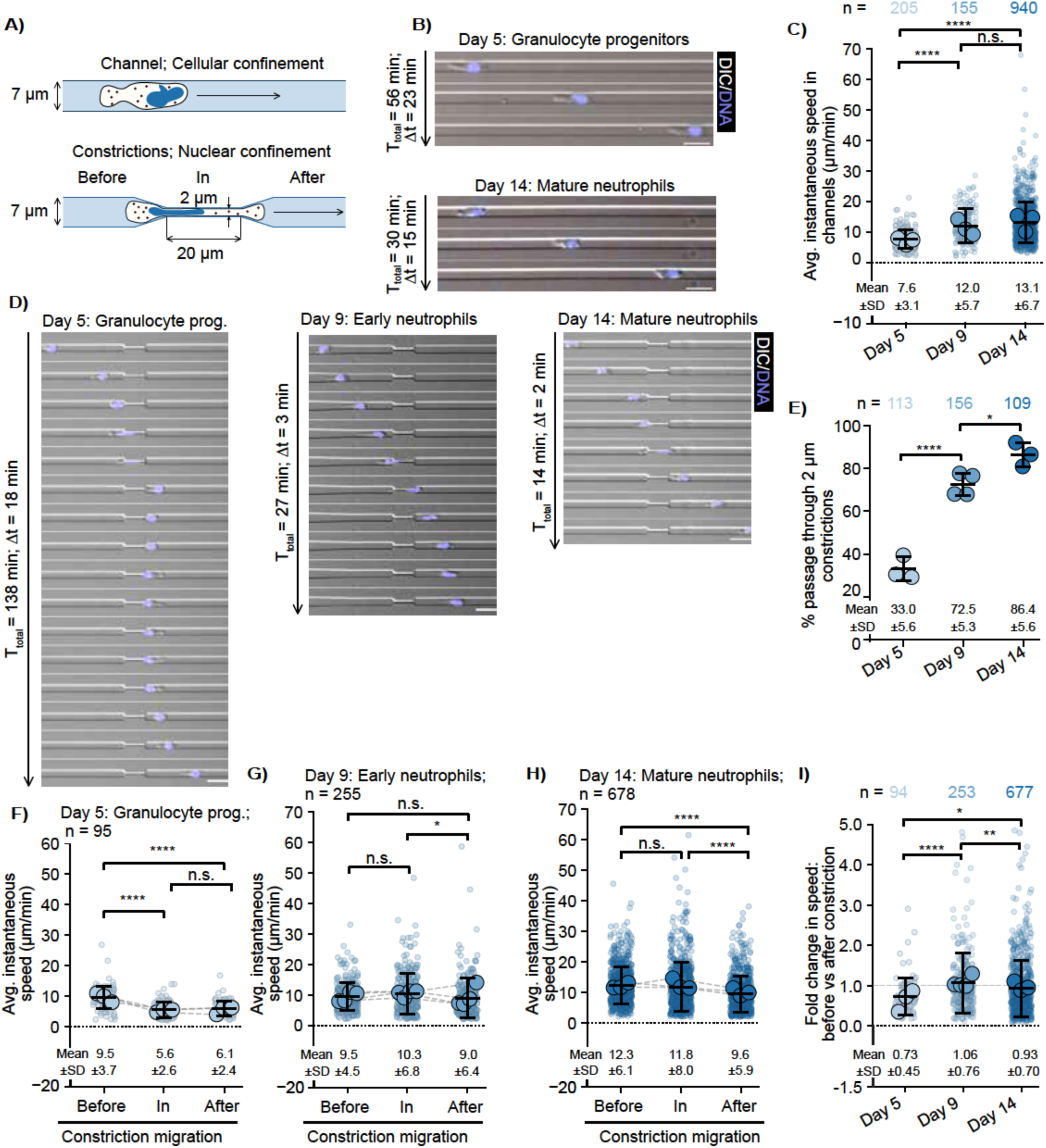
Nuclear deformation-limited migration is optimized early during neutrophil differentiation. **(A)** Schematic of the 5 μm height microfluidic devices: 7 μm wide straight channels imposing cellular confinement (top), and 7 μm wide channels containing 2 μm wide by 20 μm long constrictions imposing nuclear confinement (bottom), defining the Before, In and After regions. **(B)** Representative montages of Day 5 (Total time elapsed (T_total_) = 56 min, Time interval (Δt) = 23 min) and Day 14 (T_total_ = 30 min, Δt = 15 min) cells migrating in straight channels without constrictions. Images show overlays of DIC (grayscale) and DNA (SPY650-DNA, blue). **(C)** Average instantaneous speed of Day 5, Day 9 and Day 14 cells migrating in straight channels without constrictions. **(D)** Representative montages of Day 5 (T_total_ = 138 min, Δt = 18 min), Day 9 (T_total_ = 27 min, Δt = 3 min) and Day 14 (T_total_ = 14 min, Δt = 2 min) cells migrating in channels with 2 μm constrictions. **(E)** Percentage of cells passing through 2 μm constrictions. Each data point represents a biological replicate; n = total number of migration events. **(F - H)** Average instantaneous speed of Day 5 **(F)**, Day 9 **(G)** and Day 14 **(H)** cells in the Before, In and After regions of the constriction, measured by tracking the fluorescently labeled nucleus. **(I)** Fold change in instantaneous speed of Day 5, Day 9 and Day 14 cells, calculated per cell as the speed After divided by the speed Before the constriction; n = number of cells that fully passed through the constriction. In **(C)**, **(F - H)** and **(I)**, n = total number of cells and each data point indicates an individual cell. Bars indicate mean ± SD, shown below the graphs. Statistical tests: Mann-Whitney on 2 compared groups using pooled single-cell data in **(C)** and **(I)**; Permutation test with Bonferroni correction on 2 compared groups on cell-level paired Before/In/After triplets in **(F - H)**; pairwise Fisher’s exact tests with Bonferroni correction for three comparisons on pooled passage events and total attempts per Day in **(E)**. Data in **(C)** and **(F - I)** are shown as Superplots, where each small dot represents a cell and each large dot represents the mean of a biological replicate. n.s.: not significant; *: p-value < 0.05; **: p-value < 0.01; ***: p-value < 0.001; ****: p-value < 0.0001. Scale bars (B, D): 20 μm.

To determine when migration through constriction is optimized, we observed and analyzed granulocyte progenitors, early and mature neutrophils migrating in channels with 2 µm constrictions (**Figure 5A-bottom, D, Movie S6-S10**). Comparison of cell instantaneous speed between before and during migration in constrictions showed that granulocyte progenitors significantly decreased their speed by ∼1.7-fold in constrictions, compared to before constrictions (**Figure 5D, F**). The same analysis showed that early and mature neutrophils did not significantly decrease their speed in constrictions (**Figure 5D, G, H**). Analysis of the percentage of cells that passed through constrictions showed that ∼33%, ∼72% and ∼86% of granulocyte progenitors, early and mature neutrophils that entered, passed constrictions (**Figure 5E**). This data indicates that mature neutrophils most persistently migrate through constrictions smaller than the nucleus compared to granulocyte progenitors and early neutrophils.

We next sought to evaluate the recovery after migration through constrictions of granulocyte progenitors, early and mature neutrophils. Population level comparison of cell speed during, in and after constrictions showed that granulocytes maintain a low migration speed after constrictions and do not recover their pre-constriction speed (**Figure 5F**); early neutrophils slightly decreased their speed and recover their pre-constriction speed (**Figure 5F**); and mature neutrophils significantly decrease their migration speed and do not recover their pre-constriction speed (**Figure 5H**). Consistently, analysis of the distribution of fold change in speed between before and after constriction for each cell revealed a 0.73 ± 0.45; 1.06 ± 0.76 and 0.93 ± 0.70-fold change for granulocyte progenitors, early and mature neutrophils, respectively (**Figure 5I**). This data show that the presence of constrictions decreases granulocyte progenitors and mature neutrophils speed, in and after constriction; and that early neutrophils recover the best after migration through narrow pores.

Thus, two out of the three important features of optimal migration through physically restrictive environments (rapid migration, recovery after migration through narrow pores) are acquired early during neutrophil differentiation, while constriction-passage efficiency continues to improve with maturation.

## Discussion

In this study, we adapted a cytokine-guided in vitro differentiation pipeline to generate millions of mature human HSC-derived neutrophils and showed that rapid migration and recovery after passage through narrow pores emerges early during neutrophil differentiation. We showed that HSC-neutrophils better recapitulate the surface marker and whole proteome of blood-derived primary human neutrophils compared to HL60 derived neutrophils, the most commonly used in vitro model for human neutrophils. We showed that HSC-neutrophils phenotypically recapitulate primary neutrophils; they produce reactive oxygen species upon PMA stimulation and release neutrophil extracellular traps upon ionomycin and LPS stimulation, with comparable efficiency and using the same cellular mechanisms as primary neutrophils. Having access to human neutrophils at distinct stages of differentiation, we performed comparative proteomics and showed that the granulocyte progenitors to neutrophil transition is characterized by a proteome signature associated with decreased translation and increased immune response while the early to mature neutrophil proteome transition associates with increase in actin-regulated processes. Combining comparative proteomics and quantitative microscopy, we also showed that nuclear envelope composition is dynamically tuned during neutrophil differentiation through changes in protein expression and perinuclear localization. Notably, LBR and lamin A/C gradually increase and decrease, respectively as neutrophils differentiate while perinuclear lamin B1 and B2 abruptly decrease at the early-mature neutrophil transition. Critically, perinuclear LBR, lamin A/C, lamin B1/B2 only weakly correlate with nuclear circularity in single cells, suggesting that an ensemble nuclear envelope state regulates nuclear multilobulation. Finally, using microfluidic devices, we showed that rapid migration is optimized in early neutrophils and that while mature neutrophils are most efficient at crossing constrictions, early neutrophils recover the fastest from such migration.

Our proteome analysis of primary neutrophils, HSC neutrophils at different stages of differentiation as well as non-differentiated and neutrophil-differentiated HL60 cells, provides a resource for the field to build on and improve our understanding of neutrophil biology. HSCs are genetically tractable (48–50), allowing the use of genetic engineering to modulate protein composition for mechanistic studies of neutrophil states and functions. Our finding that HSC-neutrophils recapitulate mature human neutrophil surface markers and whole proteome indicates that our differentiation protocol produces mature neutrophils, overcoming a limitation of prior neutrophil differentiation pipelines (24). Two major findings of our comparative proteomics are that the granulocyte progenitors-early neutrophil transition is characterized by downregulation of the translation machinery and upregulation of immune response-related processes (1); while the early to mature neutrophil transition is characterized by upregulation of actin-related processes (2). Downregulation of the translation machinery supports previous work suggesting that de novo protein production is repressed in neutrophils (31, 32); we expand beyond these studies by showing that this occurs early during neutrophil differentiation indicating that it is a feature of neutrophils that is not specific to mature neutrophils. Upregulation of immune processes at the early stage of differentiation suggests that early neutrophils are immune competent. Upregulation of actin-related processes between early and mature neutrophils suggests that cellular processes regulated by the actin cytoskeleton, including migration and phagocytosis, might be optimized in mature neutrophils.

Our migration analysis temporally distinguishes the emergence of three features of optimal migration through confining spaces: rapid migration, efficient deformation through narrow pores and rapid recovery from such migration (**Figure 5**). Strikingly, our finding that rapid migration is optimized in early neutrophils while efficient passage through narrow pores is continuously optimized during neutrophil differentiation opens the door for untangling how neutrophils navigate the different physical barriers they encounter in tissue-like microenvironments (3). Our proteomics and microscopy-based analysis of nuclear shape and envelope composition showed that these parameters are tuned the most between granulocyte progenitors and early neutrophils (**Figure 4**), consistent with drastic change in constriction migration behaviors between these two differentiation stages. But our correlation analysis identified only weak correlation between nuclear circularity and perinuclear lamin A/C or LBR (**Figure 4J, K**); two proteins that gradually change during differentiation. This challenges the current view in the field that lamin A/C or LBR expression define nuclear multilobulation in neutrophils (3, 6, 51, 52). Our identification of TMPO, LEMD2, LEMD3 (**Figure S6**) as part of nuclear envelope structural proteins specifically downregulated in early neutrophils, where nuclear multilobulation emerges and rapid migration is optimized, opens the door for future investigation of their roles in regulating nuclear shape and neutrophil constricted migration. Plus, our finding that nuclear pore complex related proteins are persistently downregulated while ESCRT-related proteins are persistently upregulated raises the intriguing possibility that neutrophils modulate nucleo-cytoplasmic transport and nuclear membrane repair to potentially optimize nuclear volume regulation as well as nuclear envelope repair; two processes important for efficient nuclear deformation through narrow pores and subsequent cell survival (3, 53).

Our finding that recovery from constriction migration differs between granulocyte progenitors, early and mature neutrophils provides an opportunity to understand the emergence and regulation of cellular mechanical memory in granulocytes and neutrophils. Our data indicate that granulocyte progenitors are the most impacted by constrictions, immediately and after constrictions; followed by mature neutrophils (**Figure 5I**). Strikingly, constrictions slightly, although non-significantly, accelerate early neutrophils, and these cells resume their pre-constriction speed best. These observations raise several important questions whose answers will materially advance our understanding of granulocytes and neutrophil migration: How is this slow versus rapid recovery from constriction migration regulated in neutrophils and their progenitors? Does rapid recovery impact early neutrophils including their survival and immune functions? Is early neutrophils’ rapid recovery from constriction migration maintained in the viscoelastic tissue microenvironment?

### Limitation of the study

Our work should be evaluated within certain limitations. First, HSC neutrophils are in vitro differentiated using a cytokine cocktail that mimics aspects of emergency granulopoiesis (54, 55). Thus, our resulting HSC-neutrophils are not biological replacement of primary neutrophils. Consistently, they separate from primary neutrophils in PCA analysis (**Figure 1F**). Additionally, the high CD11b expression and low CD66b and CD10 expression of HSC-neutrophils, relative to primary neutrophils might suggest priming or activation along with lower maturation level. Although functional assays suggest that respiratory burst and NETosis of HSC-neutrophils are comparable to that of primary neutrophils, further characterization is needed to assess how well HSC-neutrophils recapitulate primary neutrophils. Second, our study aimed at characterizing neutrophils at different stages of differentiation to determine when they acquire optimal migration behaviors; a transition critical for mature neutrophil trafficking as well as the over recruitment of neutrophil progenitors in disease. Future mechanistic studies will be important to identify the regulators of neutrophil optimal migration and test their role in neutrophil immune functions. Our proteome wide characterization of these cells provides a window for such mechanistic studies.

## Materials and Methods

### Cell culture and differentiation

#### Hematopoietic stem cell expansion culture

Human bone marrow CD34+ cells (Stemcell Technologies, 70002.1) were thawed rapidly in a 37 °C water bath (<2 min) and diluted dropwise into pre-warmed HSC thawing medium (RPMI 1640 (Cytiva, SH30096.01), 10% heat-inactivated fetal bovine serum (FBS; R&D Systems, S11150)). Cells were pelleted at 300 × g for 10 min at room temperature and washed once in warm thawing medium. The pellet was resuspended at 1 × 10⁴ cells/mL in HSC expansion media (StemSpan SFEM II (Stemcell Technologies, 9655), 10% StemSpan CD34+ Expansion Supplement (10X; Stemcell Technologies, 02691) and either UM729 (Stemcell Technologies, 72332) or StemSpan HSC Plus Supplement (Stemcell Technologies, 100-1694), then plated at 1 mL per well in 24-well tissue culture–treated plates and cultured at 37 °C and 5% CO₂. Fresh pre-warmed expansion medium was added on day 4. For cryopreservation, cells were pelleted and resuspended in ice-cold freezing medium (90% heat-inactivated FBS, 10% dimethyl sulfoxide (DMSO; Sigma-Aldrich, D2650)), cooled at −80 °C for 24 h in a controlled-rate freezing container, and transferred to liquid nitrogen for long-term storage.

#### Differentiation of HSC-Neutrophils from HSC progenitors

Expanded HSCs were thawed the same as during the initiation of expansion culture and were differentiated using a three-step cytokine-driven workflow using StemSpan SFEM II (Stemcell Technologies, 9655) as basal media. For Step 1 (days 0–4), cells were cultured at 1 × 10⁵ cells/mL in medium supplemented with SCF (Stemcell Technologies, 78062), FLT3L (Stemcell Technologies, 78009.1), IL-3 (Stemcell Technologies, 78040.1), GM-CSF (Stemcell Technologies, 78015.1) and G-CSF (Stemcell Technologies, 78012). For Step 2 (days 4–7), cells were transitioned to medium supplemented with SCF and G-CSF only. For Step 3 (days 7–14), cells were transitioned to medium supplemented with G-CSF alone. Cultures were maintained at 37 °C and 5% CO₂ throughout.

Recombinant cytokines were reconstituted in sterile water to ≥0.1 mg/mL and stabilized with bovine serum albumin (BSA) to a final concentration of 0.1% (Albumin, Bovine Fraction V; Research Products International, A30075). Stocks were aliquoted and stored at −80 °C. Cytokines were used at final concentrations of 50 ng/mL for SCF, FLT3L, IL-3 and GM-CSF and 100 ng/mL for G-CSF.

#### HL60 cell culture and differentiation

A human promyelocytic leukemia cell line, HL60 cells, was purchased from ATCC (ATCC® CCL-240™), cultured in RPMI 1640 medium (Cytiva, SH30096.01) supplemented with 1% glutamax (Thermo Scientific, 35050061), 1% Penicillin/Streptomycin (Pen/Strep; Gibco, 15070063), 25 mM HEPES (Cytiva, SH30237.01) and 15% heat-inactivated Fetal Bovine Serum (FBS; R&D Systems, S11150) and split every 3 days at 2×10^5^ cells/mL. Cells were maintained at 37°C in a humidified 5% CO_2_ incubator. Cells were differentiated into dHL60 cells via treatment with culture media supplemented with 1.3% dimethyl sulfoxide (DMSO; Sigma-Aldrich D2650). Differentiated cells were used at day 6 or 7 after treatment with DMSO.

#### Isolation of peripheral blood neutrophils

Human blood samples were obtained from healthy donors from the Stanford Blood Center. Research blood donors provided written informed consent and blood samples were de-identified prior to distribution. Neutrophil isolation was performed using a commercially available immune-magnetic negative selection kit (EasySep™ Direct Human Neutrophil Isolation Kit, Stemcell, 19666). Whole blood was mixed with magnetic beads and negative-selection antibody cocktail before being placed into ‘The Big Easy’ EasySep™ Magnet (Stemcell, 18001) to remove all non-neutrophil cells in the whole blood. Cells were washed 1x with HBSS prior to usage for experiments.

### Flow cytometry for surface marker expression

Cells were resuspended in 4% PFA in Hank’s Balanced Salt Solution (HBSS, Sigma-Aldrich, H9394), fixed for 20 min at 37 °C, and quenched in 0.1 M glycine in HBSS for ≥5 min. Cells were washed twice in ice-cold FACS buffer (BD Biosciences, 554656). For antibody staining, 1 × 10⁶ cells were resuspended in 100 µL FACS buffer and single-stained using 5 µL of fluorophore-conjugated antibody: CD10 BV421 (BD Biosciences, 562902), CD66b PE (BD Biosciences, 561650), CD15 APC (BD Biosciences, 561716), CD16 R718 (BD Biosciences, 566970), and CD11b BB515 (BD Biosciences, 564517). Cells were incubated in the dark for 30 min at 4°C, then washed twice in FACS buffer and analyzed on a Sony SH800 cell sorter using a 100 µm sorting chip with standard instrument calibration and chip alignment using calibration beads. Gates were defined using unstained controls and single-stained controls for each fluorophore, with channel-specific boundaries set by peak shifts relative to unstained cells. Samples were run on medium flow rate and detection was stopped at 100,000 cells. Percentage of cells that were positive for surface-marker expression was measured with BD FlowJo v10.

### Proteomics sample preparation and data acquisition

Abundance proteomics samples from cell pellets were prepared using the SPEED method (Sample Preparation by Easy Extraction and Digestion) (56). 8x10^5^ cells were collected for (1) freshly isolated primary human neutrophils, (2) HL60 and dHL60 cells at day 6 of differentiation, and (3) HSC-Neutrophils at days 5, 9, and 14 of neutrophil differentiation. All samples were prepared in biological triplicates except primary human neutrophils, which were prepared in biological quadruplicates (cells derived from four different donors). Following two washes with phosphate-buffered saline (PBS) for 5 min at 400 × *g,* cell pellets were lysed with 10 μL of 100% trifluoroacetic acid (TFA, Fisher Chemical, A116-50) and incubated for 2 minutes, lysates were then neutralized with 2M Tris Base at 10x the volume of TFA. Extracted proteins were reduced and alkylated using 10 mM Tris (2-carboxyethyl) phosphine hydrochloride (TCEP, Sigma Aldrich, C4706-10G) and 40 mM Chloroacetic acid (CAA, Sigma Aldrich, C19627-25G) for 5 minutes at 95°C. Protein concentrations of the lysates were determined using Pierce 660 protein assay (Thermo Fisher, 22660). 10 µg total protein from each lysate was diluted 1:2 using H2O and then digested at 37°C for 20 hours at 800rpm using Trypsin (Promega, V5113) and LysC (Fujifilm Biosciences NC9309054) at an enzyme to protein ratio of 1:50. Following digestion, peptides were acidified, desalted, dried, and resuspended in 25 µL 0.1% formic acid (FA, Thermo Scientific, 85178). 1 µL of peptide suspension (∼400 ng total protein) was injected for liquid chromatography-mass spectrometry analysis. Samples were analyzed using a TimsTOF HT mass spectrometer (Bruker) coupled to a Vanquish Neo ultra high-pressure liquid chromatography system (Thermo Fisher Scientific) via a CaptiveSpray2 nanoelectrospray source. Peptides were loaded onto an IonOpticks 15 cm x 75 μm I.D. Aurora CSI column. Mobile phase A consisted of 0.1% FA, and mobile phase B consisted of 0.1% FA/80% acetonitrile (ACN). Peptides were separated at a flow rate of 500 nL/min using a nonlinear gradient increasing buffer B from 2% to 80% over 53-minutes followed by a set 7-minute column wash and equilibration. All MS experiments acquired data in dia-PASEF mode using a 75 ms TIMS ramp and utilized a TIMS precursor MS window covering an *m/z* range of 100-1700 Da and a 1/K_0_ range of 0.72-1.50. One TIMS precursor MS frame was followed by 9 dia-PASEF MS2 frames using fragmentation windows having a width of 30 Da with an m/z overlap of 1 Da between windows. Fragmentation windows spanned a m/z range of 250-1325 Da and a 1/K0 range of 0.72-1.50.

### Proteomics data analysis

All resulting RAW files were analyzed with Spectronaut (Biognosys, v20.5) using direct DIA analysis for the identification and quantification of the resulting dia-PASEF and synchro-PASEF datasets (57). The following changes to the default BGS factory settings were made for all processed data. Cross-run normalization was turned off. Acquired data were searched against the UniProt canonical human proteome (downloaded October, 2024) and the BGS contaminant database (57, 58). Carbamidomethylation of cysteine (C) was used as a fixed modification, and protein N-terminus acetylation and methionine oxidation were set as variable modifications. The PSM (peptide spectral matches), peptide, and protein group false discovery rates were set to 0.01 with no missing value imputation enabled. Following pre-processing the resulting peptide fragment ion intensities were exported from Spectronaut in *tsv* format and were imported into R (v 4.5.1) for further analysis using the MSstats framework (59). After importing transition-level data from Spectronaut into the R environment, peptides from known contaminant proteins were filtered. Data were converted to the standard MSstats format using the *SpectronauttoMSstatsFormat* function using both a protein - and peptide-level qvalue_cutoff of 0.01. These data were summarized to protein-level abundances using the MSstats *dataProcess* function (4.16.1). Only features marked as ‘highQuality’ were retained. Data were normalized using the *equalizeMedians* argument and missing value imputation was disabled. For Figure 1, Principal component analysis was done using the 3420 proteins shared across all replicated and all conditions. Next, the *groupComparison* function was used to run pairwise comparisons of quantified proteins between the experimental conditions versus control. Thresholds of q ≤ 0.05 (Benjamini-Hochberg correction); |log_2_ FC| ≥ 1 were used to determine significantly changing proteins compared to control. Gene Ontology enrichment analysis was conducted using the R package clusterProfiler (v.4.14.4) and associated enricher function (60). Biological processes were retrieved directly from Bioconductor’s org.Hs.eg.db database. Redundancy in GO terms was minimized by incorporating the simplify function (61). For sample-level analyses, protein intensities were reshaped to a protein-by-sample matrix and correlation matrices calculated from protein-level log intensities. For figure 3, hierarchical clustering was performed on the z-score-transformed abundance matrix using Euclidean distance and complete linkage, and proteins were partitioned into discrete temporal clusters using cutree. Cluster-level trajectories were summarized by calculating the mean z-score across all proteins within a cluster for each sample and then averaging within each condition. For figure 4, a curated protein list was generated from the Human Protein Atlas (47) (available from: v25.proteinatlas.org. Accessed July 16, 2026) using search results for “nuclear envelope proteins”. Proteins present in both the Human Protein Atlas-derived list and the proteomics dataset were retained. As in Figure 3, proteins filtered to those showing significant differential abundance in at least one pairwise comparison between D5, D9, and D14 cells (adjusted *p* < 0.05). Protein abundances were row-wise z-score transformed and hierarchically clustered, and cluster-level temporal trajectories were calculated as described above. Nuclear envelope proteins were submitted to STRING for protein–protein interaction analysis using a high-confidence interaction score threshold. Networks were imported into Cytoscape for visualization, with nodes colored according to their temporal z-score abundance trajectories. All code used for analysis is present at this GitHub link.

### Intracellular Reactive Oxygen Species Detection

Intracellular reactive oxygen species (ROS) production was measured using a DCFDA/H2DCFDA Cellular ROS Assay Kit (Abcam, ab113851). Cells were resuspended at 1 × 10⁶ cells/mL in a total volume of 1 mL and washed once with room-temperature Dulbecco’s phosphate-buffered saline (DPBS). Cells were then resuspended in phenol red-free RPMI 1640 medium supplemented with 10% heat-inactivated FBS and DCFDA dye at a 1:1000 dilution and incubated for 30 min at 37 °C in the dark. After staining, cells were washed once with pre-warmed HL60 imaging medium and seeded at 1 × 10⁵ cells in 100 µL per well in a 96-well plate. PMA-containing medium was added to each stimulated condition to achieve a final PMA concentration of 20 nM, and cells were incubated for an additional 30 min at 37 °C. Fluorescence was measured on a CLARIOstar Plus multimode microplate reader (BMG Labtech) using excitation and emission wavelengths of 485 and 535 nm, respectively, according to the manufacturer’s recommendations. Two replicate wells were analyzed per condition. Following plate-reader analysis, cells were imaged using a 20× objective on an ECHO Revolve microscope (RVL2-K3).

### Immunostaining

#### Cell fixation and permeabilization

5 × 10⁴ HSC-Neutrophils in 5 µL imaging medium were seeded onto plasma-cleaned (Harrick Plasma, PDC-001) 12 mm #1.5 glass coverslips (Electron Microscopy Sciences, 72290-04) and allowed to adhere for 5–10 min at 37 °C. Cells were fixed in 4% paraformaldehyde (Electron Microscopy Sciences, 15710) prepared in 1× cytoskeleton buffer (CB; 10 mM MES, 138 mM KCl, 3 mM MgCl₂, 2 mM EGTA) for 20 min at 37 °C. Cells were permeabilized with 0.5% Triton X-100 in 1× CB for 5 min at 37 °C, and unreacted PFA was subsequently quenched in 0.1 M glycine in 1× CB for at least 5 min. Coverslips were washed twice (5 min each) at room temperature with washing buffer (0.1% Tween-20 (Sigma-Aldrich, P7949) in 1× Tris-buffered saline (TBS; 20 mM Tris-HCl pH 7.6, 150 mM NaCl)).

#### Indirect immunofluorescence

Fixed and permeabilized cells were blocked in 2% BSA and 0.1% Tween-20 in 1× TBS for 1 h at room temperature. Primary antibodies were diluted in blocking solution and incubated with coverslips overnight at 4°c. Coverslips were washed three times (5 min each) in washing buffer (0.1% Tween-20 in 1× TBS), then incubated for 1 h at room temperature with fluorophore-conjugated secondary antibodies diluted in blocking solution together with DAPI (1 µg/mL; Sigma-Aldrich, D9542) and Alexa Fluor 647 Phalloidin (Cell Signaling Technologies, 8940S). Coverslips were washed three times (5 min each) in washing buffer and mounted in fluorescence mounting medium (Dako, S302380-2).

Primary antibodies used were rabbit anti-LBR (Abcam, ab32535; 1:1,000), mouse anti-lamin A/C (Abcam, ab232730; 1:1,000), mouse anti-lamin B2 (Abcam, ab8983; 1:1,000), mouse anti-lamin B1 (Abcam, ab8982; 1:1,000), and rabbit anti-PAD4 (GeneTex, GTX113945; 1:1,000). Secondary antibodies used were goat anti-rabbit Alexa Fluor 488 (Invitrogen, A-11008; 1:1,000) and goat anti-mouse Alexa Fluor 568 (Invitrogen, A-11031; 1:1,000).

### Microfluidic device fabrication

#### Photolithographic mold production

Photomasks were designed in AutoCAD 2025 and custom-ordered as chrome transparency photomasks. Silicon wafers (University Wafers) were rinsed with acetone, methanol, and isopropanol, then dried for 10 min at 95°C. SU-8 2005 negative photoresist (Kayaku Advanced Materials, Y111045) was spin-coated and exposed on a mask aligner (Carl Suss) to form a 5 µm-thick adhesion layer. A second 5 µm SU-8 2005 layer was subsequently spin-coated and exposed through the photomask to create the device features for a device with an indented 5 µm height. Unexposed SU-8 was washed away in SU-8 developer (Kayaku Advanced Materials, Y020100), rinsed with isopropanol to stop development, and hard-baked for 2 h at 120°C.

To prevent PDMS adhesion, wafers with master molds were vapor-silanized with trichloro(1H,1H,2H,2H-perfluorooctyl)silane (Sigma-Aldrich, 448931). A sacrificial PDMS layer (Sylgard 184 Silicone Elastomer Kit; Sigma-Aldrich, 761036) mixed at a 10:1 (base:curing agent, w/w) ratio was cast onto the silanized master and cured at 80°C for 2 h, then peeled to remove non-covalently associated silane and condition the mold surface for subsequent device fabrication. Device heights were verified using a contact profilometer.

#### PDMS device fabrication

Sylgard 184 was mixed in a 10:1 (base:curing agent, w/w) ratio and cast on the wafer. After 1 hour of degassing in a vacuum desiccant, the wafer was baked at 80°C overnight. PDMS layer was gently peeled off and inlet holes for cells were punched using a 2 mm tissue biopsy punch. PDMS device molds and #1.5 glass bottom 6-well plates (Cellvis; P06-1.5H-N) were air-plasma treated for 1 min before bonding and baked at 80°C for at least 15 minutes to complete the PDMS bonding.

#### PDMS device functionalization and cell seeding

Devices bonded to glass bottom 6-well plates were placed in the plasma cleaner with the lid off and vacuum was applied for 10 minutes before plasma cleaning for 60 seconds. After venting the chamber, a P200 pipette was used to dispense 50 ug/mL human fibronectin protein (EMD Millipore, FC010) in Dulbecco’s Phosphate Buffered Saline (DPBS, ThermoFisher, 14190144) into each well. Devices were placed in 37°C + 5% CO_2_ incubator for 1 hour. Afterwards, devices were washed in imaging media (serum-free, no phenol red RPMI 1640 medium (Gibco, 11835030) buffered with 25 mM HEPES, 1% Pen/Strep, 1x Glutamax) and left immersed in imaging media in an incubator at 37°C and 5% CO_2_ for at least 2 hours. Cells were stained for 1 hour at 37°C and 5% CO_2_ in StemSpan SFEM II media in a well of a 24 well plate with a fluorescent DNA dye (1:1000, SPY650-DNA, Cytoskeleton, CY-SC501) before being washed 2x in imaging media and being seeded at density of 100,000 cells per 5 μL per well of the microfluidic device. Devices were placed for 1 hour at 37°C and 5% CO_2_ before being covered in imaging media. Devices were placed for at least 1 more hour in the incubator before being taken for live imaging.

### Microscopy

#### Spinning disc confocal and DIC microscopy

Imaging was performed on a Nikon Eclipse Ti2 inverted microscope equipped with Perfect Focus, an Okolab stage-top incubator for controlled temperature, humidity and CO_2_, a Crest X-light V3 spinning disc scanhead, a Kinetix sCMOS camera, a Nikon LUNF (405nm, 90mW) and Nikon Opti-Microscan-01 for FRAP/Photostimulation, and the appropriate DIC prisms in place. Illumination was provided on the Crest V3 by a 7-line solid-state Celesta Laser Unit, 1 W per channel (405 nm; 445 nm; 488 nm; 515 nm; 561 nm; 640 nm; and 730 nm power measurement is at the fiber output). DIC illumination was provided by an LED. Microscope was equipped with the Nikon motorized stage with xy linear encoders and a closed-loop *Z*-axis Piezo nanopositioning system with 200 μm travel. Laser confocal or DIC illumination was selected with electronic shutters and an automated filter turret containing a multibandpass dichromatic mirror together with an electronic emission filterwheel. Microscope functions were controlled with NIS-Elements software (Nikon).

#### Live cells NETosis assays

Cells were seeded into #1.5 glass bottom 12 well plates (Cellvis, P12-1.5H-N). Random fields (3-10) were selected with adhered cells. Fluorescence confocal, DIC or brightfield images were acquired using a Plan Apo 40× air 0.95 NA DIC Nikon objective lens at the coverslip-cell interface for each position. Images were acquired every 60 seconds for 4 hours at maximum 5% laser power (for 640 nm) and 2x2 binning. 5 minutes from the start of the movie, NETosis stimulants were applied: 20 µg/mL Lipopolysaccharides from Klebsiella (Sigma-Aldrich, L4268) or 4 µM Ionomycin (Sigma-Aldrich, I0634).

#### Live cells migration in microfluidic devices

4-15 fields of views were chosen based on channel regions closest to the well where cells were seeded. Fluorescence confocal, DIC or brightfield images were acquired using a Plan Apo 20× air 0.8 NA DIC Nikon objective lens at the coverslip-cell interface for each position. Images were acquired every 30 or 60 seconds for 8 hours at maximum 5% laser power (for 640 nm laser) and 2x2 binning.

#### Fixed cell imaging

Fixed and stained cells mounted on slides were imaged through the coverslip using a Plan Apo 60× oil 1.4 NA DIC Nikon objective lens. Random fields (4 to 6) containing >50 adherent cells were selected, and pairs of DIC and fluorescence confocal stacks were captured. Images were acquired with 405 nm, 488 nm, 561 nm and 647 nm at 50% laser power and no binning.

### Image Analysis

#### Quantification of the percentage and timing of cellular events during NETosis

Time-lapse movies of cells stained with SPY650-DNA were used to quantify the percentage and timing of NETosis events as previously described (15, 62). Events quantified: Plasma membrane microvesicle shedding, defined as the release of vesicles from the cell periphery as seen in DIC movies; chromatin reorganization, defined as the decrease in SPY650-DNA heterogeneity as seen in confocal fluorescence movies; Nuclear rounding, defined as the establishment of a circular nuclear periphery as seen in DIC and confocal fluorescence color overlay movies; Nuclear rupture, defined as expansion of the DNA periphery outside of the nuclear boundaries into the cytosol as seen in DIC and confocal fluorescence color overlay movies; PM permeabilization, defined as a decrease in contrast at the cell periphery as seen in DIC movies; and NETosis, defined as expansion of DNA outside of the cell boundary as seen in DIC and confocal fluorescence color overlay movies. All adherent cells in imaging fields were included in the analysis of the percentage of cells undergoing an event while only cells that remained in the field for the entire duration of the movie were included in the quantification of the timing of events. Multiple imaging fields were analyzed for each condition until the number of cells per condition was equal or greater than 50. The timing of an event was defined as the first time point when the corresponding event was observed in the movies by eye.

#### Quantification of migration speed

Segmentation and tracking code was adapted from Kang, et al (63). Images from time-lapse movies were first converted into binary masks and segmented on a frame-by-frame basis using an automated watershed-based pipeline that includes Gaussian smoothing, distance-transform marker generation, and morphological hole filling, followed by exclusion of objects outside an area range to remove debris and aggregates. For each segmented object, centroid coordinates were extracted and converted from pixels to physical units using a calibrated scale factor, and a per-frame table of object features was assembled. Individual cells were then linked across consecutive frames using a global assignment (Hungarian) approach that minimizes a cost function based on squared centroid displacement with an additional penalty for large frame-to-frame size changes, yielding continuous cell tracks over time. Instantaneous migration speed was calculated from centroid displacements between successive timepoints divided by the imaging interval, and tracks were partitioned into “before”, “In”, and “after” constriction segments based on the cell’s position relative to the constriction region along the channel axis; segment-specific speeds were computed by aggregating instantaneous speeds within each region. Cells were tracked for 200 µm before and after the constriction.

#### Quantification of nuclear passage through constrictions

For each experiment, up to 20 constrictions directly parallel to the entry well of the device were selected for analysis. Within each field of view, up to 50 total migration events were scored manually across the duration of the movie. A migration event was defined as a cell entering the constriction region; successful passage was scored when the cell fully traversed the constriction (In) and exited to the post-constriction channel (After). For each independent biological variable, the percentage of passage was calculated by pooling the total number of successful passages across all scored positions and dividing by the total number of migration events attempted across those same positions.

#### Quantification of protein of interest (POI) and nuclear shape in fixed cells

Fixed-cell images were acquired as 16-bit multi-channel, multi-Z stacks on a spinning disk confocal microscope. Fixed-cell immunofluorescence images were analyzed in Python (NumPy, scikit-image, SciPy) using a custom script applied to single-cell-cropped TIFF stacks. For quantification, z-stacks were collapsed using an average-intensity projection. To quantify nuclear protein enrichment, the projected DAPI channel was used to generate a nuclear mask by automated thresholding (Triangle), conversion to a binary mask, and hole-filling, followed by erosion. Background was estimated as the mean intensity outside this mask and subtracted from each channel prior to quantification. The DAPI-based mask was then multiplied by the corresponding protein channel to restrict signal to the nucleus, with integrated intensities measured within the masked region; mean nuclear intensities were calculated by normalizing integrated intensity by the number of nuclear pixels. Nuclear shape features (e.g. circularity) were instead derived from the LBR channel, independent of the DAPI mask used for intensity quantification. The best-focus Z-plane of the LBR channel was selected per cell by Laplacian-variance focus scoring; LBR pixel intensities within the DAPI-defined nuclear region were thresholded at a fixed percentile cutoff (10th percentile, retaining the top ∼90% of signal), holes were filled, the largest connected region was retained, and the resulting mask was Gaussian-smoothed and re-thresholded to yield the final nuclear outline used for shape quantification. Samples lacking an LBR stain were quantified for nuclear protein/DNA intensity as above; LBR intensity and all shape metrics were not computed for these samples. Pixel-to-micron conversion was performed using the imaging calibration (µm per pixel) specified at analysis time, and all measurements were exported per cell to a single results table for downstream statistics.

### Statistical analysis and reproducibility

#### Study design and reproducibility

All experiments were performed in N ≥ 3 independent biological replicates, defined as independent differentiations initiated from separate CD34+ HSC lots (Stemcell Technologies, 70002.1) derived from at least five different bone marrow donors. Primary human neutrophils were isolated from independent healthy blood donors. For single-cell measurements we aimed for n > 50 cells per condition to minimize the standard error of the mean, which is proportional to one over the square root of n. Imaging fields were selected randomly without reference to cell phenotype. Blinding was not relevant to most analyses, which were automated; manual scoring (NETosis event timing, constriction passage) was performed without prior knowledge of the expected phenotype. Micrographs and montages are representative of the cell populations quantified within the same figure. The differentiation protocol was independently executed and validated by five experimenters at different career stages (postbaccalaureate researcher, graduate student and postdoctoral fellow).

#### Data exclusion

No data were excluded from the flow cytometry, ROS or fixed-cell analyses. In the migration analysis, exclusion criteria were built into the analysis pipeline and applied identically across all conditions and timepoints: (i) segmented objects below 50 px² were discarded as debris; (ii) frame-to-frame links were rejected when centroid displacement exceeded 200 px or when object area changed by more than 3.5-fold; (iii) region-specific speeds were computed only for cells that traversed all three regions, so cells reaching only the “Before” region, or only “Before” and “In”, were retained for the passage-rate analysis but excluded from the speed analysis; and (iv) tracks were truncated when a cell re-entered the constriction after reaching the “After” region, and frames following five or more consecutive frames with an unchanged centroid in the “After” region were excluded as stationary. For nuclear passage analysis, scoring was capped at 50 total migration events per biological replicate across up to 20 constrictions parallel to the entry wall. For the proteomics analysis, one biological replicate (D9.2) was excluded from downstream analysis due to a documented contamination event during differentiation, occurring independently of sample collection or MS processing; raw data for this replicate were retained but excluded from all reported comparisons and analyses. In the NETosis analysis, cells that migrated out of the field of view where excluded from analyses.

#### Statistical methodology

Analysis was performed in Python v3.12.11 (SciPy v1.16.2; NumPy v2.3.3; pandas v2.3.3) with figures generated in matplotlib v3.10.6; proteomics analysis was performed in R v4.5.1 as described above. Data are presented as mean ± SD on figures below the graph data. Lowercase *n* denotes individual cells or migration events, and uppercase *N* denotes independent biological replicates; the applicable definition is stated in each figure legend. Distributions of fluorescence intensity, nuclear circularity and migration speed were non-normal, and non-parametric tests were used throughout if values were not log-adjusted: Mann– Whitney U for two-group comparisons; Permutation test with Bonferroni correction for paired “Before”/“In”/“After” comparisons within the same cells; Fisher’s exact test with Bonferroni connection was applied to the pooled counts across three pairwise day comparisons for comparing percentage of cellular passage through constrictions. Ratio comparisons of DMSO- versus PMA-treated wells were performed using a paired *t*-test on log-transformed values, equivalent to a one-sample *t*-test on the log fold change. Proportions (percentage of cells reaching each NETosis stage, percentage of cells passing through constrictions) were compared using Fisher’s tests on the underlying counts and were not analyzed as continuous variables. Spearman’s rank correlation coefficient was used to assess monotonic associations between nuclear circularity and nuclear envelope protein levels, as these relationships were expected to reflect progressive, stage-dependent change rather than a constant rate of change. Where tests were performed on cells pooled across replicates, data are displayed as Superplots (64) so that replicate-to-replicate variation is visible alongside the pooled comparison. P-values greater than 0.05 were considered non-significant and are not annotated; * denotes p < 0.05, ** p < 0.01, *** p < 0.001 and **** p < 0.0001.

## Data availability statement

Proteomics data has been deposited to the ProteomeXchange Consortium via the MassIVE partner repository with the dataset identifier PXD083298 and MSV000103033 respectively (65). All other data are available from the corresponding authors upon request.

## Code availability

All analysis codes are available on this Github link.

## Acknowledgements

We thank the funding agencies who support our work. A.Y is supported by Stanford Bioengineering and the NIH BTP training grant (5T32GM141819). K.O. is supported by the Stanford Biosciences and NIGMS MCB training grant (T32GM154663). H.R.T. and the Thiam lab are supported by the Biohub, San Francisco, the David and Lucile Packard Foundation, Stanford Bio-X, the Koret Foundation, and the Esther Ehrman Lazard Faculty Scholar Award. We thank Alejandro Matià for help with the proteomics samples preparation and Dain Brademan for help with the proteomics data analysis. We thank members of the Thiam lab for constructive feedback on the project, figures and manuscript.

## Author contributions

A.Y., K.O., R.S.F., and H.R.T. conceptualized the study. A.Y. and K.O. conducted the study, and prepared figures. R.S.F. developed the initial HSCs differentiation protocol and performed initial characterization of nuclear envelope composition and functional NETosis assays. A.D. performed differentiation and immunostaining assays. R.H. designed and supervised the proteomics experiments. H.R.T designed experiments, supervised the work and acquired funding for the study. A.Y., K.O and H.R.T. wrote the manuscript. All authors read and corrected the article and contributed to the interpretation of results.

## Competing Interest

The authors declare no competing interests.

## Supplementary Information

**Supplemental Figure 1:**
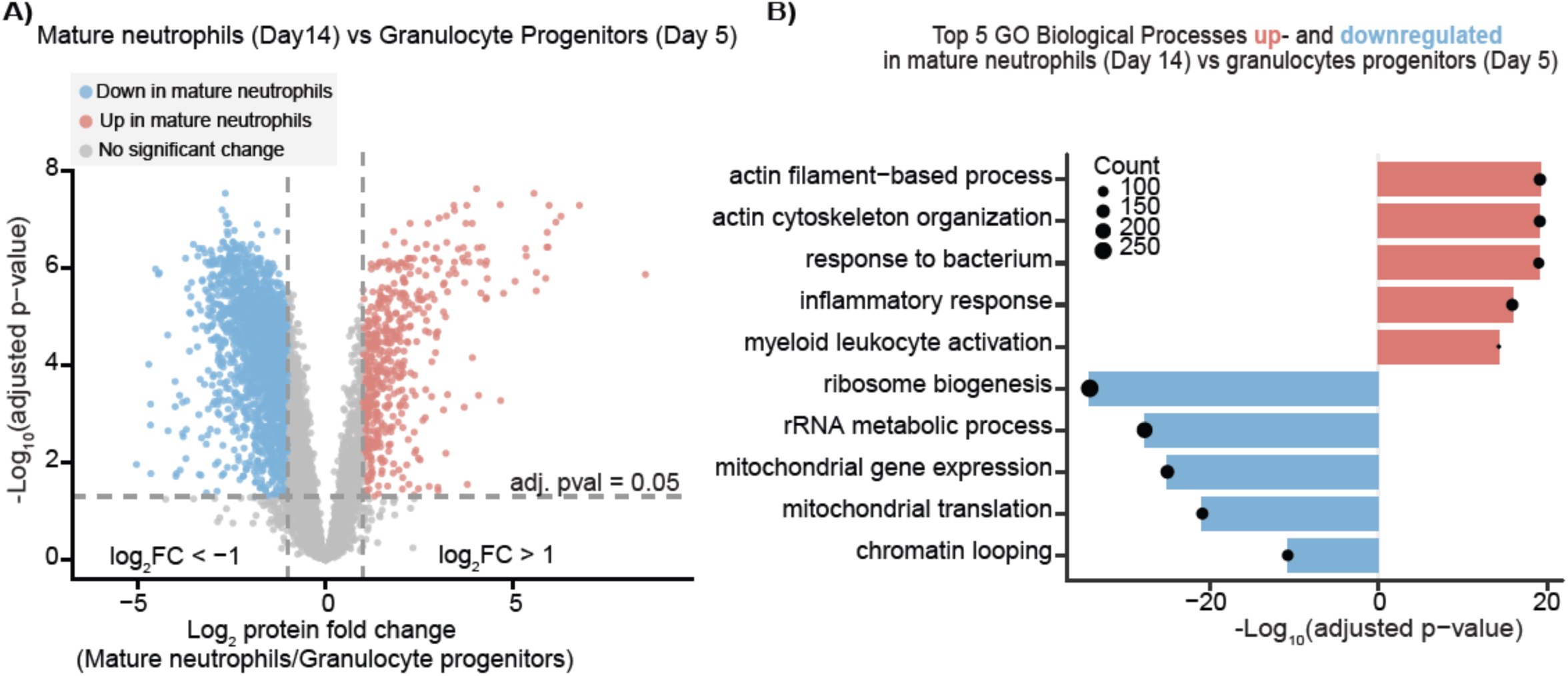
HSC-Neutrophils maintain high viability for 7 days after differentiation. **(A)** Cell viability (%) of HSC-neutrophils measured daily from Day 14 to Day 21 after the onset of differentiation. Each dot represents a biological replicate, and Each line connects the daily measurements of one biological replicate; N = 3 independent biological replicates / differentiations. The pink band marks Day 18, on which media exchange was performed. Mean ± SD are shown below the graph.

**Supplemental Figure 2:**
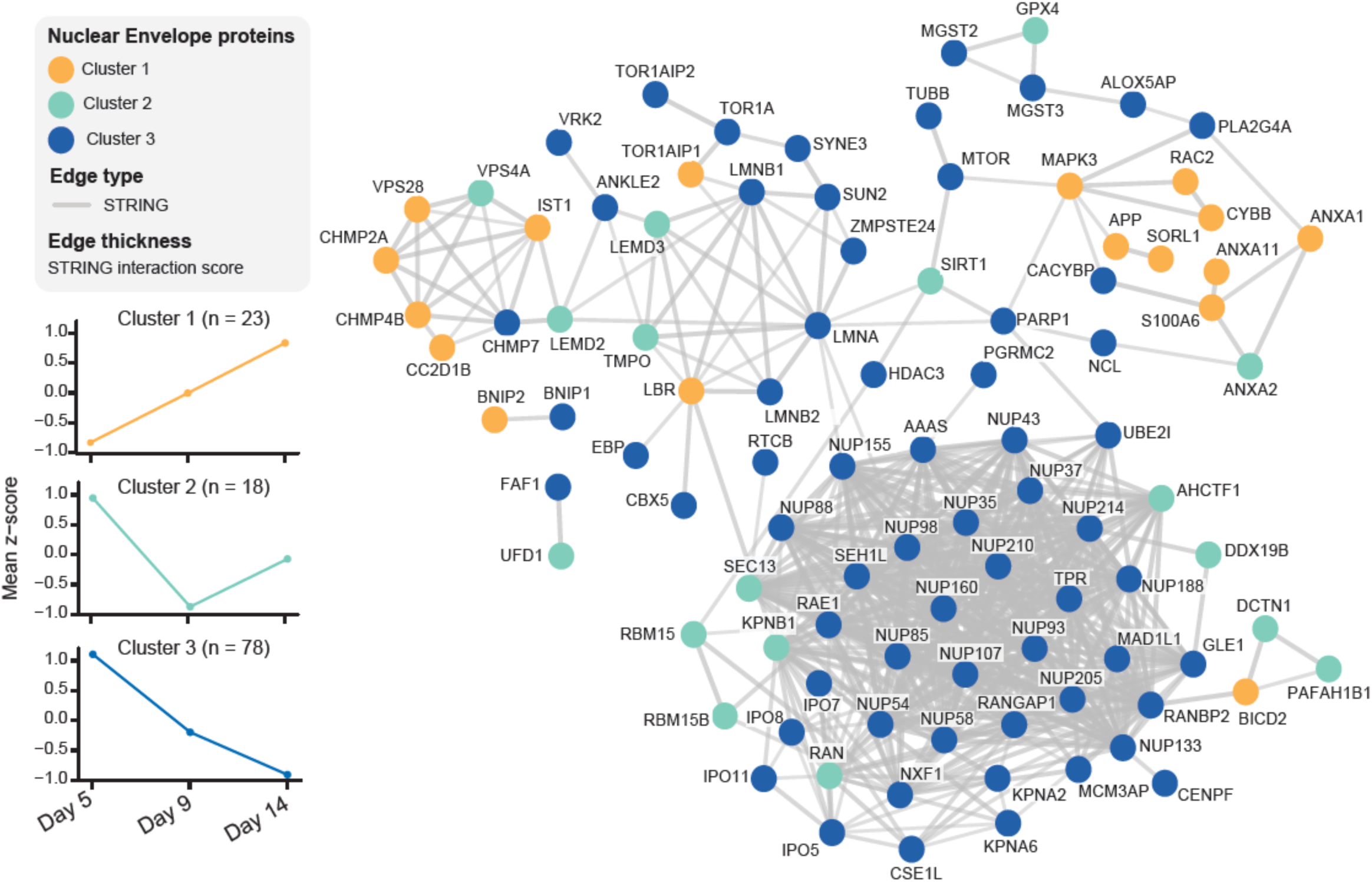
dHL60 cells do not fully recapitulate primary human neutrophil surface markers. **(A**) Representative flow cytometry histograms of CD16, CD15, CD66b, CD10 and CD11b signal in unstained primary neutrophils (gray), stained primary neutrophils (red) and stained dHL60 cells (orange). Dashed line indicates the positivity gate set on the unstained primary neutrophil control. Histograms are representative of N ≥ 3 biological replicates, quantified in Figure 1D.

**Supplemental Figure 3:**
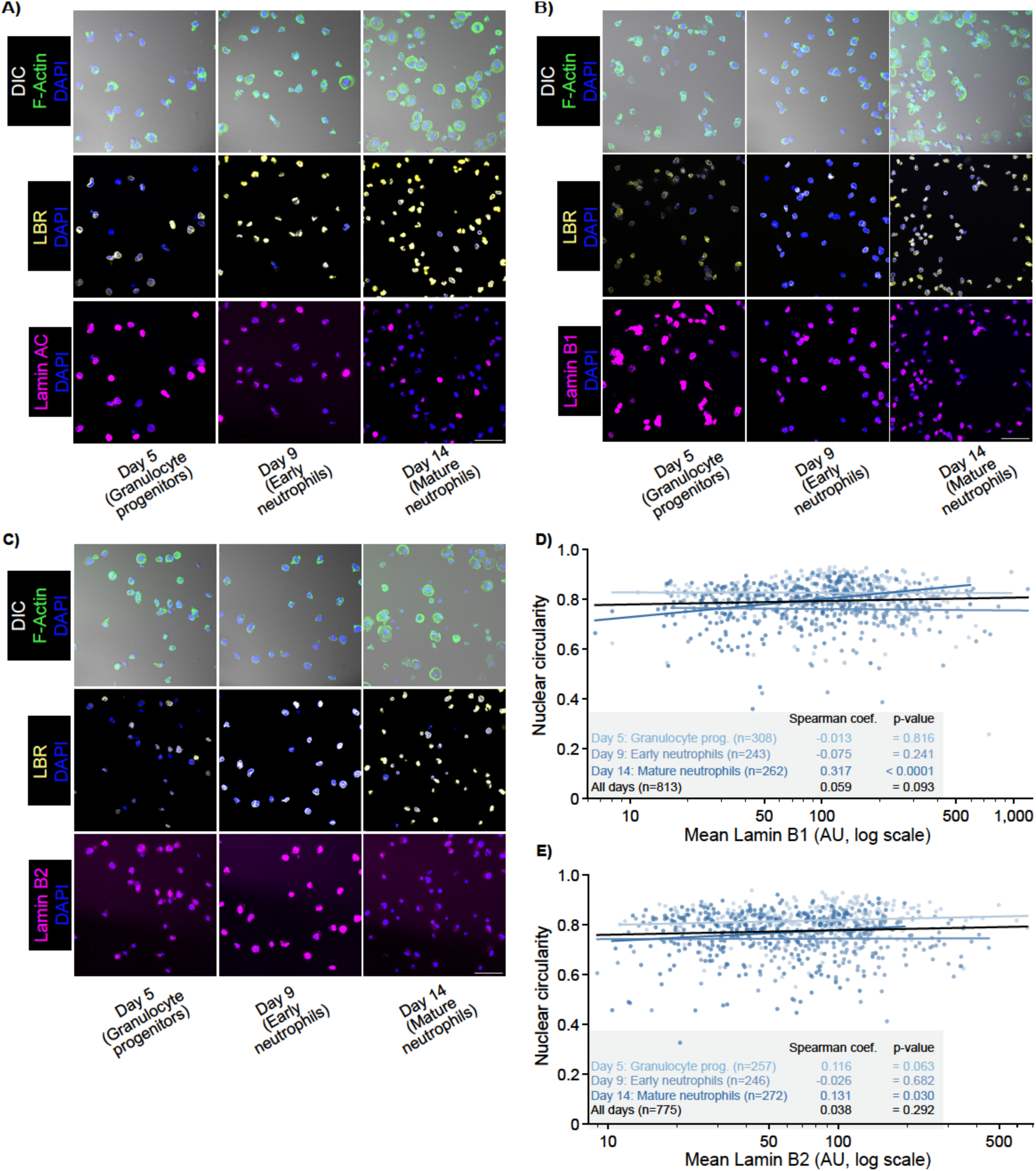
Quantitative proteomics shows that HSC-neutrophils better recapitulate primary neutrophil proteome when compared to dHL60 cells. **(A)** Number of identified peptide sequences for each replicate for cells indicated on the x-axis, with respective mean and standard deviation (SD). **(B)** Number and distribution of protein groups identified in each replicate for cells indicated on the x-axis, with respective mean and SD. **(C)** Number of unique protein groups identified per cell type across replicates. **(D - E)** Spearman correlation analysis of the mean log_10_ intensities between primary neutrophils and HSC-neutrophils (n = 5384 proteins, **D**) and between primary neutrophils and dHL60 cells (n = 5307 proteins, **E**). **(F)** Spearman correlation analysis matrix of all cell populations, (Day 5, Day 9 and Day 14 HSC-neutrophils, primary neutrophils, HL60 and dHL60) showing replicate correlations averaged into a single box. Heat to visualize spearman coefficient from 0 < r_s_ <1. **(G)** Mean log₂ intensities of neutrophil maturation markers: c-kit/CD117 (hematopoietic progenitors/immature myeloid cells marker), FUT4 (CD15-producing enzyme marking myeloid commitment), CD66b/CEACAM8 (granulocyte maturation marker), CD11b/ITGAM (adhesion marker), CD16/FCGR3B (late stage, mature neutrophil marker), and CD10/MME (mature neutrophil marker)).

**Figure S4:**
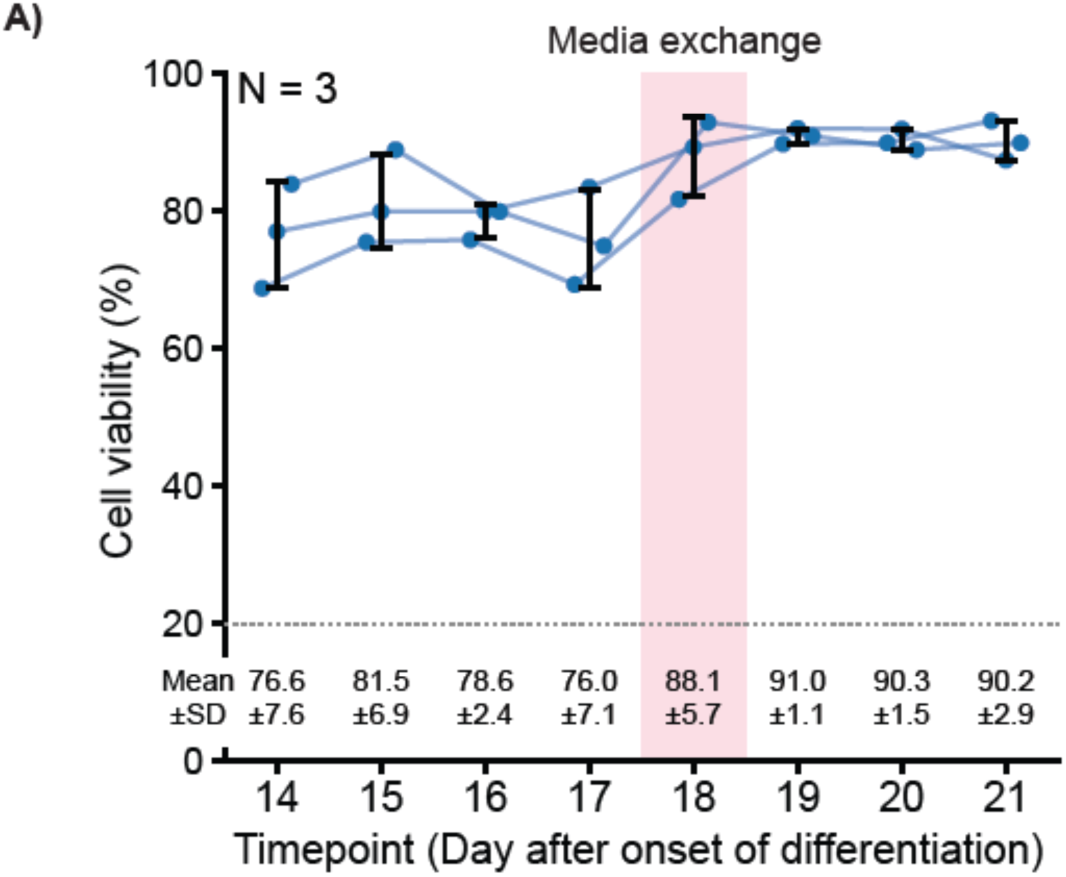
Primary and HSC-Neutrophils have minimal DCF background fluorescence. **(A)** Representative images of primary neutrophils (top) and HSC-neutrophils (bottom) that were not loaded with DCFDA and were treated with DMSO (Control) or PMA (20 nM) under the same conditions as in Figure 2A. Left: images displayed with the brightness and contrast settings used in Figure 2A. Right: the same images displayed with increased contrast so that individual cells are visible. The unstained primary human neutrophil condition was performed in N = 3 independent experiments and is quantified in Figure 2B; the unstained HSC-neutrophil condition was performed once (N = 1) and is not quantified in Figure 2B. Scale bar: 40 μm.

**Figure S5:**
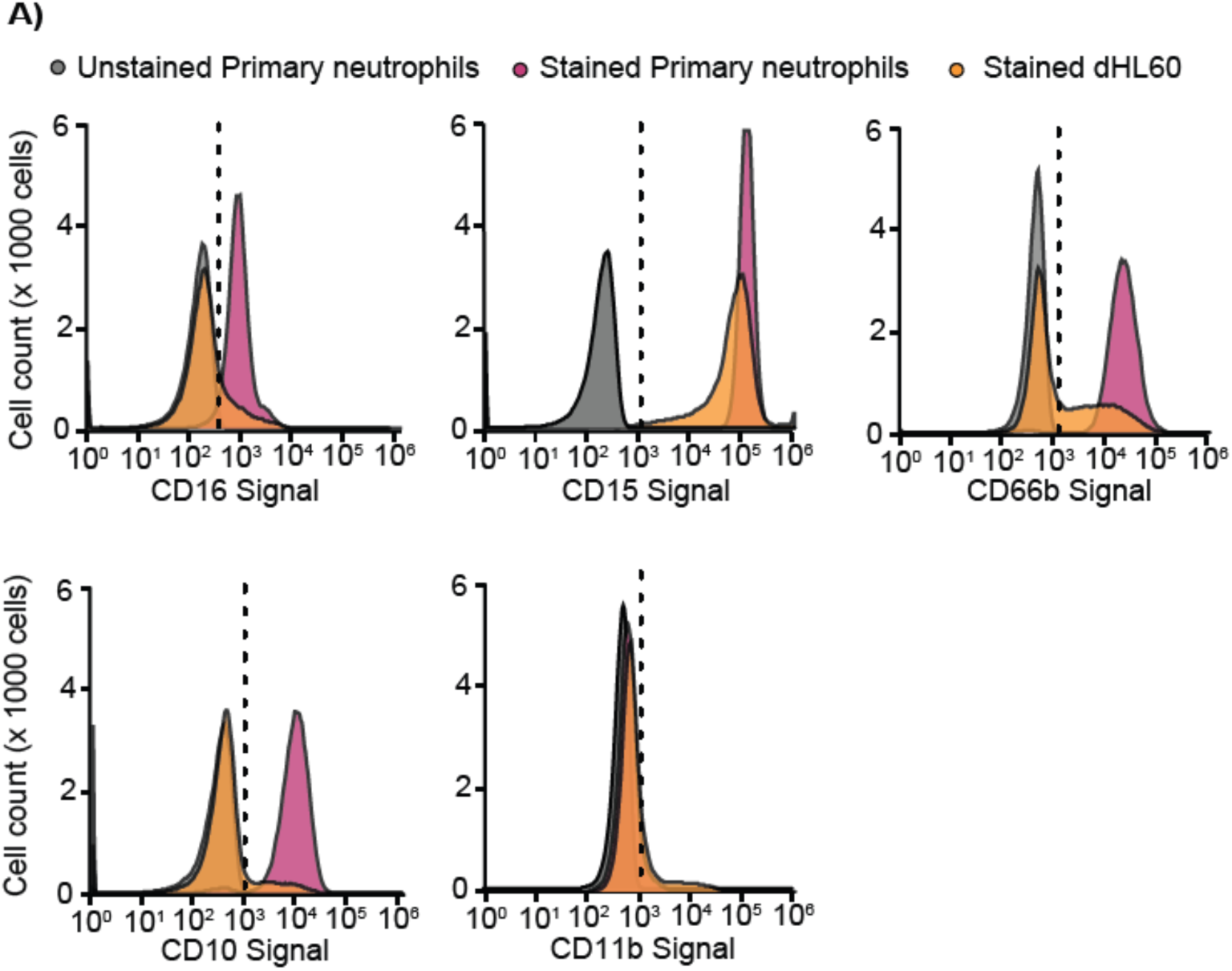
Single cell dynamics of Primary and HSC-Neutrophils undergoing NETosis upon ionomycin stimulation. **(A)** Representative montages of primary (top) and HSC-neutrophils (bottom) stimulated for NETosis with Ionomycin (4 μM) and quantified in (Figure 2G-I). Images show overlays of DIC (grayscale) and SPY650-DNA (blue) at the indicated stages of NETosis: non-stimulated, NETosis onset, chromatin reorganization, nuclear rounding, nuclear rupture and cell rupture.

**Figure S6:**
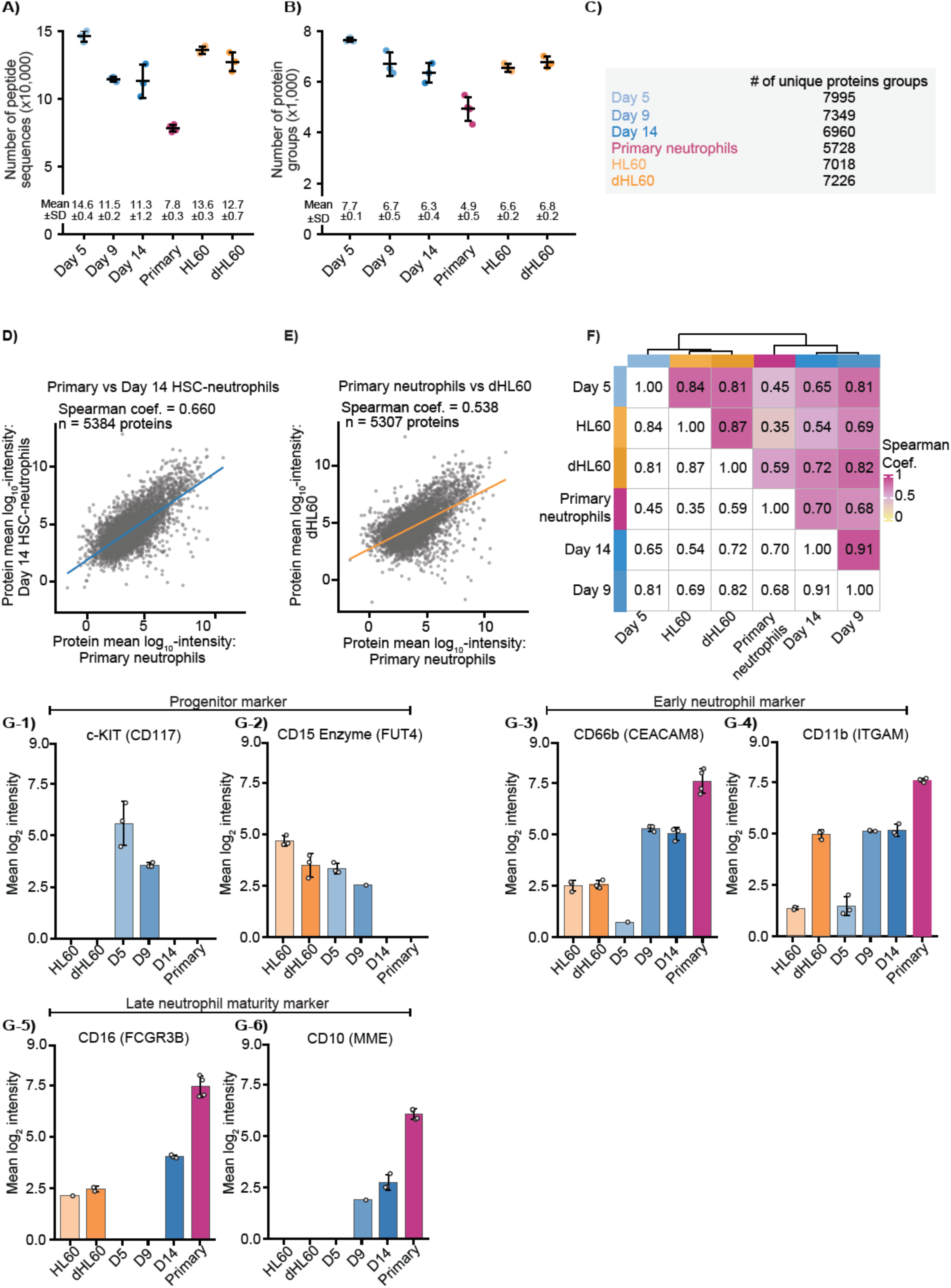
HSC neutrophils gain immune response and actin organization and become transcriptionally repressed relative to granulocyte progenitors. **(A)** Volcano plot showing differential protein abundance for Mature neutrophils vs Granulocyte progenitors (|log₂FC| ≥ 1, adjusted p-value ≤ 0.05) **(B)** GO enrichment analysis showing the top 5 most enriched biological processes within the upregulated (red) and downregulated (blue) proteins in mature neutrophils compared to granulocyte progenitors.

**Figure S7:**
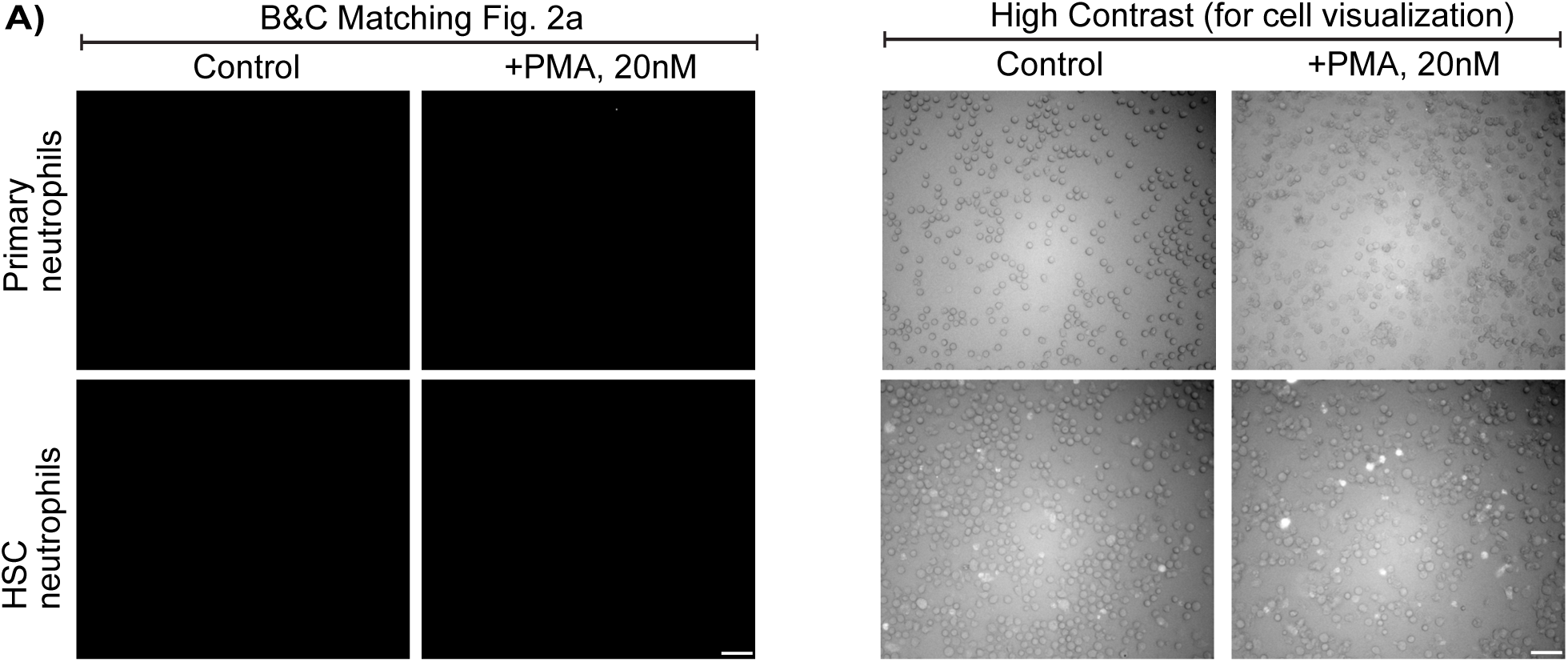
Nuclear envelope proteins are dynamically remodeled during neutrophil differentiation. **(A)** STRING interaction network of differentially expressed nuclear envelope proteins from (Figure 4A) colored by their cluster. Z-score trajectories of each cluster are shown in corresponding colors.

**Figure S8:**
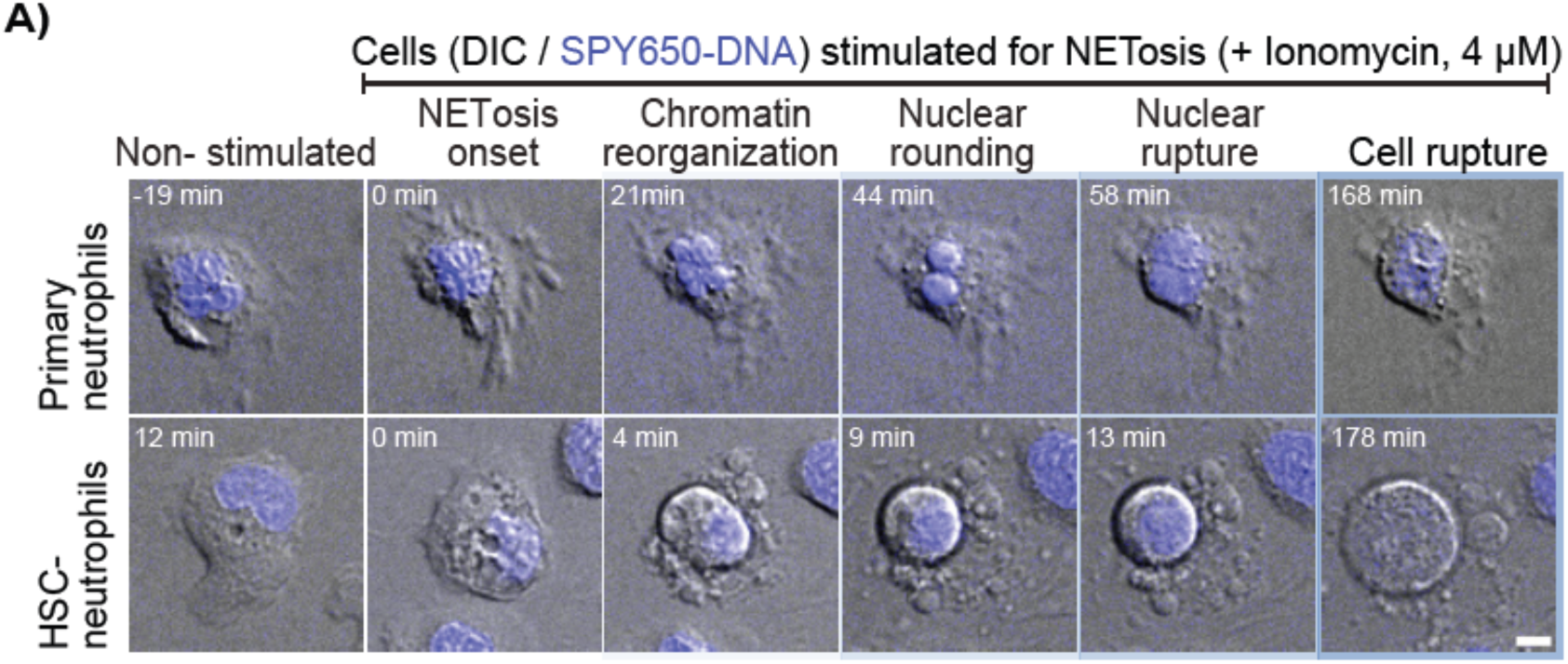
Nuclear multilobulation increases during HSC to neutrophil differentiation but does not correlate with perinuclear lamin B1 or B2 signal in all cells. **(A - C)** Representative large field of view (200 × 200 μm regions) images of Day 5, Day 9 and Day 14 cells corresponding to the single-cell images in Figure 4B, D and E and quantified in Figure 4 D, G-I. Cells stained for LBR (yellow), DAPI (blue) and lamin A/C (magenta, **A**), lamin B1 (magenta, **B**) or lamin B2 (magenta, **C**). Top: DIC overlaid with F-actin (green) and DNA (DAPI, blue). Middle: LBR (yellow), overlaid with DNA (DAPI, blue); Bottom: lamin A/C, B1 or B2 (magenta), overlaid with DNA (DAPI, blue). **(D, E)** Single-cell correlation analysis between nuclear circularity and mean perinuclear lamin B1 (**D**) or mean perinuclear lamin B2 (**E**) intensities; x-axis on a log scale. Lines indicate best fit for each day (color codes in blue) and for all days pooled (black); Spearman coefficients and associated p-values are reported in the inset table. Fluorescence intensities in **(D)** and **(E)** were measured as the mean pixel intensity per nucleus within a nuclear mask defined by the DAPI signal and are expressed in arbitrary units (AU). Bars indicate mean ± SD, shown below the graph. Scale bars: 40 μm.

Supplemental Movies Legends: Link to Google Drive

**Movie S1: Day 14 HSC-neutrophil and primary neutrophil undergoing NETosis following LPS stimulation.**

Side-by-side view of Day 14 HSC-neutrophil (left) and primary neutrophil (right), both stained with SPY650-DNA, undergoing NETosis following stimulation with 20 μg/mL LPS. Overlay of DIC (grayscale) and SPY650-DNA (blue). Images were acquired at 1 min intervals for 4 hours. Scale bar: 5 μm. Playback: 10 fps.

**Movie S2: Day 14 HSC-neutrophil and primary neutrophil undergoing NETosis following ionomycin stimulation.**

Side-by-side view of Day 14 HSC-neutrophil (left) and primary neutrophil (right), both stained with SPY650-DNA, undergoing NETosis following stimulation with 4 μM ionomycin. Overlay of DIC (grayscale) and SPY650-DNA (blue). Images were acquired at 1 min intervals for 4 hours. Scale bar: 5 μm. Playback: 10 fps.

**Movie S3: Day 5 granulocyte progenitors migrating in a straight channel.**

Day 5 cells stained with SPY650-DNA migrating in a 7 μm wide straight channel under cellular confinement. Overlay of DIC (grayscale) and SPY650-DNA (blue). Images were acquired at 1 min intervals. Scale bar: 20 μm. Playback: 10 fps.

**Movie S4: Day 14 mature neutrophils migrating in a straight channel.**

Day 14 cells stained with SPY650-DNA migrating in a 7 μm wide straight channel under cellular confinement. Overlay of DIC (grayscale) and SPY650-DNA (blue). Images were acquired at 1 min intervals. Scale bar: 20 μm. Playback: 10 fps.

**Movie S5: Day 5 granulocyte progenitor migrating through a 2 μm constriction.**

Day 5 cell stained with SPY650-DNA migrating through a single 2 μm wide by 20 μm long constriction under nuclear confinement. Overlay of DIC (grayscale) and SPY650-DNA (blue). Images were acquired at 1 min intervals. Scale bar: 20 μm. Playback: 10 fps.

**Movie S6: Day 9 early neutrophil migrating through a 2 μm constriction.**

Day 9 cell stained with SPY650-DNA migrating through a single 2 μm wide by 20 μm long constriction under nuclear confinement. Overlay of DIC (grayscale) and SPY650-DNA (blue). Images were acquired at 1 min intervals. Scale bar: 20 μm. Playback: 10 fps.

**Movie S7: Day 14 mature neutrophil migrating through a 2 μm constriction.**

Day 14 cell stained with SPY650-DNA migrating through a single 2 μm wide by 20 μm long constriction under nuclear confinement. Overlay of DIC (grayscale) and SPY650-DNA (blue). Images were acquired at 1 min intervals. Scale bar: 20 μm. Playback: 10 fps.

**Movie S8: Day 5 granulocyte progenitors migrating through a series of constrictions.** Day 5 cells stained with SPY650-DNA migrating through channels containing multiple constrictions in series. Overlay of DIC (grayscale) and SPY650-DNA (blue). Images were acquired at 1 min intervals. Scale bar: 20 μm. Playback: 10 fps.

**Movie S9: Day 9 early neutrophils migrating through a series of constrictions.**

Day 9 cells stained with SPY650-DNA migrating through channels containing multiple constrictions in series. Overlay of DIC (grayscale) and SPY650-DNA (blue). Images were acquired at 1 min intervals. Scale bar: 20 μm. Playback: 10 fps.

**Movie S10: Day 14 mature neutrophils migrating through a series of constrictions.** Day 14 cells stained with SPY650-DNA migrating through channels containing multiple constrictions in series. Overlay of DIC (grayscale) and SPY650-DNA (blue). Images were acquired at 1 min intervals. Scale bar: 20 μm. Playback: 10 fps.

## Notes

### Competing Interest Statement

The authors have declared no competing interest.

